# Liquid-solid phase transitions in the biological condensates of a conserved mitotic spindle regulator

**DOI:** 10.64898/2026.04.08.717243

**Authors:** Amalia S. Parra, Christopher A. Johnston

**Affiliations:** Department of Biology, University of New Mexico, Albuquerque, NM, 87131

## Abstract

Formation of biological condensates through phase separation has emerged as a means of discrete subcellular organization permitting regulatory precision across diverse processes. We recently found that the conserved mitotic spindle positioning complex of Mushroom body defect (Mud) and Partner of Inscuteable (Pins) phase separates into condensates with dynamic, liquid-like properties. While the Mud/Pins complex functions at the cell cortex in diverse cell types, Mud also controls centrosome activity independently of Pins at spindle poles. Here we find that Mud alone undergoes homotypic phase separation, producing condensates with characteristics unique from those of the heterotypic Mud/Pins complex. Specifically, Mud condensate droplets display a less dynamic behavior with reduced propensity for liquid-like fusions. Instead, they coalesce into larger aggregates of otherwise individual droplets, resulting in liquid-to-solid phase transitions over time. Structural modeling and mutational analyses implicate self-interacting oligomerization as a possible molecular model for this condensate behavior. Phosphorylation of Mud by the mitotic kinases, Warts or Polo-like kinase (Plk1), results in highly liquid-like condensates that fail to undergo phase transition. Lastly, we describe similar yet distinct solid formations in two additional spindle pole proteins, TACC and NudE, implicating an underlying common role for coiled-coil domains in these phenomena. Our studies identify new biophysical aspects of Mud function and highlight a role for phase transitions in the biological condensates of spindle-associated coiled-coil proteins.

## INTRODUCTION

Proper assembly of the mitotic spindle is essential for the fidelity of cell division. This demands the coordinated activity of numerous microtubule-associated proteins in mitosis that control spindle geometry, length, and orientation to ensure correct chromosome segregation and proper cleavage furrowing^1^. Among these are force-generating microtubule motor proteins that organize microtubules, congress and align sister chromatids, and maintain spindle pole integrity^2^. The minus-end-directed Dynein motor plays an essential role in each of these functions, with a particular importance at spindle poles. Dynein, together with the Dynactin adaptor complex, focuses microtubule minus ends to maintain spindle pole structure and spindle bipolarity, while also executing poleward transport of additional regulatory proteins. Dynein is also anchored at the cell cortex, where it exerts pulling forces on astral microtubules to control spindle orientation^3,4^.

Mushroom body defect (Mud), as well its functional mammalian homolog the Nuclear Mitotic Apparatus (NuMA) protein, associates with the Dynein/Dynactin complex and acts as a critical regulator of its activity^5^. Mud is a large, microtubule-binding coiled-coil protein that localizes to spindle poles and the cell cortex where it influences Dynein localization and activity^6–12^. NuMA association with Dynein is well established to control spindle pole focusing during spindle assembly, an essential aspect of spindle bipolarity and proper chromosome segregation^13–16^. A similar function for Mud at spindle poles is incompletely resolved, with studies finding conflicting evidence for a role in pole focusing possibly due to cell-specific effects^9,17,18^. Mud loss has been associated with supernumerary centrosomes and improper centrosome clustering^9,17^, phenotypes that it shares following deficits in NuMA or Dynein function. For their roles in spindle orientation, Mud and NuMA polarize to discrete cortical domains where they instruct cortex-spindle forces through their association with cortical Dynein^3,7,9,12,19–21^. Cortical polarization of Mud relies on its direct binding to the Partner of Inscuteable (Pins) protein, with an analogous pattern seen between NuMA and its cortical binding partner LGN^19^. Notably, whereas loss of Pins results in defects in cortical Mud localization and spindle positioning, Mud localization and function at spindle poles appear to be independent of Pins binding^8,12^. Furthermore, cortical Mud-dependent spindle positioning at epithelial tricellular junctions does not require Pins^6,22^. Additional studies have found that, in fact, LGN binding can negatively regulate NuMA-dependent stabilization of microtubules during spindle assembly^13^. These essential yet distinct functions of Mud in spindle regulation substantiate a better understanding of its complex molecular function^23,24^, including what molecular mechanisms might distinguish between Pins-dependent and -independent activities at specific subcellular structures.

We recently discovered that the Mud/Pins complex forms biological condensates through phase separation^25^. Phase separation has emerged as a critical mode of regulation for a growing number of protein complexes across diverse biological functions^26^. Recent studies have similarly found that NuMA phase separates, with the Dynein-binding C-terminal region playing an essential role in formation of condensates that accumulate at spindle poles during mitosis^27,28^. Our previous results showed that Mud/Pins condensates are highly dynamic *in vitro*, undergoing liquid-like fusions over time that generate large, wetted surface areas^25^. Condensate formation was dependent on direct binding between Mud and Pins, as well as the multivalency generated from proper assembly of a Mud coiled-coil domain (Mud^CC^) that is just N-terminal to the Pins-binding domain (Mud^PBD^). We also identified the actin-binding protein Hu li tai shao (Hts; Adducin in mammals) as a Mud-binding protein that could form a tripartite Hts/Mud/Pins complex. Mud-dependent recruitment of Hts to condensates reduced their liquid-like behavior, demonstrating a capacity for regulation of their dynamic behavior^25^. Lastly, Hts showed cortical co-localization with Mud and Pins in asymmetrically dividing neuroblasts, consistent with its role in Mud/Pins-dependent spindle orientation rather than a function at spindle poles. Whether phase separation plays a role in Pins-independent Mud activity has not been addressed.

We find herein that Mud is capable of forming homotypic condensates in the absence of Pins or Hts. In contrast with the dynamic, liquid-like condensates formed by the Mud/Pins complex, those of Mud alone are generally resistant to droplet fusion events, instead coalescing into larger structures we refer to here as ‘droplet-of-droplets’. Over time, these reactions form gel-like solids that also appear to be composed of numerous coalesced condensate droplets. Structural modeling supports previous biochemical findings of direct self-association between the Mud^CC^ and Mud^PBD^ domains^8^, and suggest this interaction may trigger oligomeric assemblies critical to condensate behavior. Phosphomimetic mutation at a previously identified Warts kinase site within Mud^CC^, or at a Polo kinase site within Mud^PBD^ newly identified in this study, each disrupt condensate coalescence and solid transition, producing instead dynamic droplets that undergo liquid-like fusions similar to the Mud/Pins complex. Lastly, we find that two additional coiled-coil spindle pole proteins, the Transforming Acidic Coiled-Coil (TACC) protein and NudE, each also phase separate into solid structures. Our results provide new insights into the biophysical properties of critical spindle pole proteins.

## RESULTS

### Mud undergoes homotypic phase separation in vitro

We recently described the ability of the Mud/Pins complex to phase separate *in vitro*^25^. In this case, the spherical condensates produced displayed a highly dynamic behavior evident by liquid-like droplet fusions over time, ultimately leading to large, amorphous droplets with extensive surface wetting. Herein, we have identified homotypic Mud phase separation that occurs independent of Pins. Specifically, we find that a purified Mud fragment spanning a contiguous cassette of coiled-coil (CC), ‘Linker’, and PBD domains (referred to here as Mud^1955^; Figure 1A) produced extensive phase separated spherical droplets *in vitro* using a minimal buffer in the absence of crowding agents or additional additives. Droplet formation and behavior was sensitive to ionic strength and pH. When examined at a constant pH of 7.5, the number and size of droplets were observed to vary across a range of ionic strength. Specifically, a higher number of individual droplets were observed at the lower range of NaCl concentrations (50-100mM), and these droplets were variable in size (Figure 1B). A higher NaCl concentration range (200mM), fewer droplets were observed and those that did form were more uniformly of smaller size. Further increase in ionic strength (300mM) abolished condensate formation. However, the physical behavior of droplets (liquid-like dynamics) was similar across the entire range, with relatively few droplet fusions occurring over time despite their close proximity (Figure 1B). When examining condensate formation at a constant ionic strength (100mM NaCl), altering pH across a physiological range primarily affected the liquid-like dynamics of droplets. At acidic pH (6-6.5), droplets were highly dynamic, undergoing extensive liquid-like fusions that ultimately formed large amorphous droplets and a wetted surface (Figure 1C). At alkaline pH (8-8.5), droplets were highly stable as individual spherical entities and largely resisted fusion events. Neutral pH (7.5) produced an intermediate phenotype that generally favored stable droplet behavior with minimal fusions (Figure 1C). Together, these results demonstrate that the protein domains within Mud^1955^, which are unique to the specific Mud isoform known to control mitotic spindle function^12,29^, are sufficient to induce condensate formation and suggest that the extent of phase separation and the behavior of the resulting droplets are influenced by electrical charge dynamics of the Mud protein and its surrounding bulk solution.

**Figure 1.**
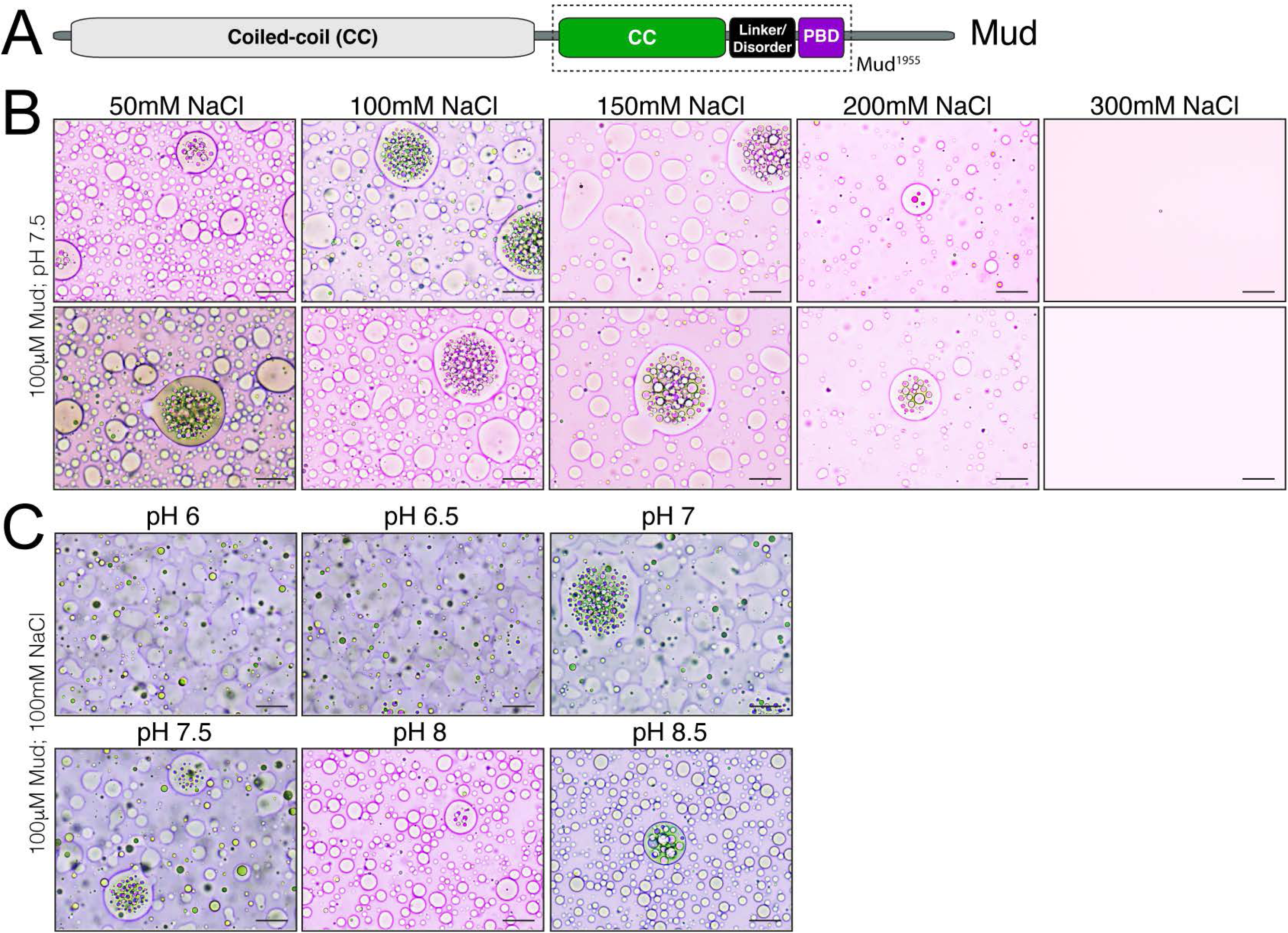
Mud forms homotypic phase-separated condensates. **(A)** Domain architecture of Mud shows it is primarily composed of coiled-coil (CC) domains. An extended N-terminal CC domain (*light grey*) is commonly present in Mud isoforms and not the focus of the current study. An additional, smaller CC domain (*green*) within a C-terminal region is separated from the Pins-binding domain (PBD; *purple*) by a disordered ‘Linker’ segment (Linker/Disorder; *black*). These three tandem domains (*dashed box*, termed Mud^1955^ herein) are encoded by isoform-specific exons and are the focus of the current study^12^. **(B)** Mud^1955^ protein was diluted to 100μM final concentration in buffer at pH 7.5 (20mM Tris) and the indicated concentrations of NaCl (columns). Images in top and bottom rows are from identical conditions of independent reactions to provide two examples of each condition. Scale bars, 10μm. **(C)** Mud^1955^ protein was diluted to 100μM final concentration in buffer with 100mM NaCl and the indicated pH (20mM HEPES for 6-6.5; 20mM Tris for 7-8.5). Scale bars, 10μm.

An additional, rather conspicuous aspect of the Mud^1955^ condensates was their apparent coalescence into ‘clusters’ of larger aggregates containing otherwise individual droplets (Figure 1). This ‘droplet-of-droplets’ phenotype was observed across the ranges of ionic strength and pH examined, suggesting it is a core aspect of Mud^1955^ condensate behavior. These clusters were always encompassed within a large droplet that itself was distinct from smaller droplets and the surrounding bulk solution, although these large droplets were sometimes found to undergo droplet fusions (Figure 1B-C). Despite liquid-like fusions between the large droplet with smaller, isolated surrounding droplets, the coalesced droplets contained within it were not observed to fuse over time despite their close proximity.

### Mud condensates undergo a liquid-to-solid phase transition

During our analysis of Mud^1955^ phase separation above, we noticed under standard conditions (100μM protein in 100mM NaCl, pH 8) the formation of a semi-translucent, gel-like solid apparent at the bottom of the reaction tube on visual inspection (Figure 2A). This solid formed after ∼2 hours of incubation, was seen at both 20°C and 4°C, and was stable for at least several weeks. This structure was resistant to disruption by gentle pipetting, allowing for an intact transfer to a microscope slide soaked in its original reaction solution (Figure 2A). Microscopic inspection revealed that the bulk reaction solution contained individual condensate droplets similar to those shown previously. The solid structure, in contrast, was densely packed with numerous individual droplets coalesced within the solid boundary (Figure 2B). This droplet coalescence was strikingly similar to the phenotype described above, albeit within a solid matter of significantly larger size. The individual droplets within the solid varied in size but were immobile and did not undergo fusion events despite their close proximity to one another. Additional imaging of these structures is shown in Figure S2A.

**Figure 2.**
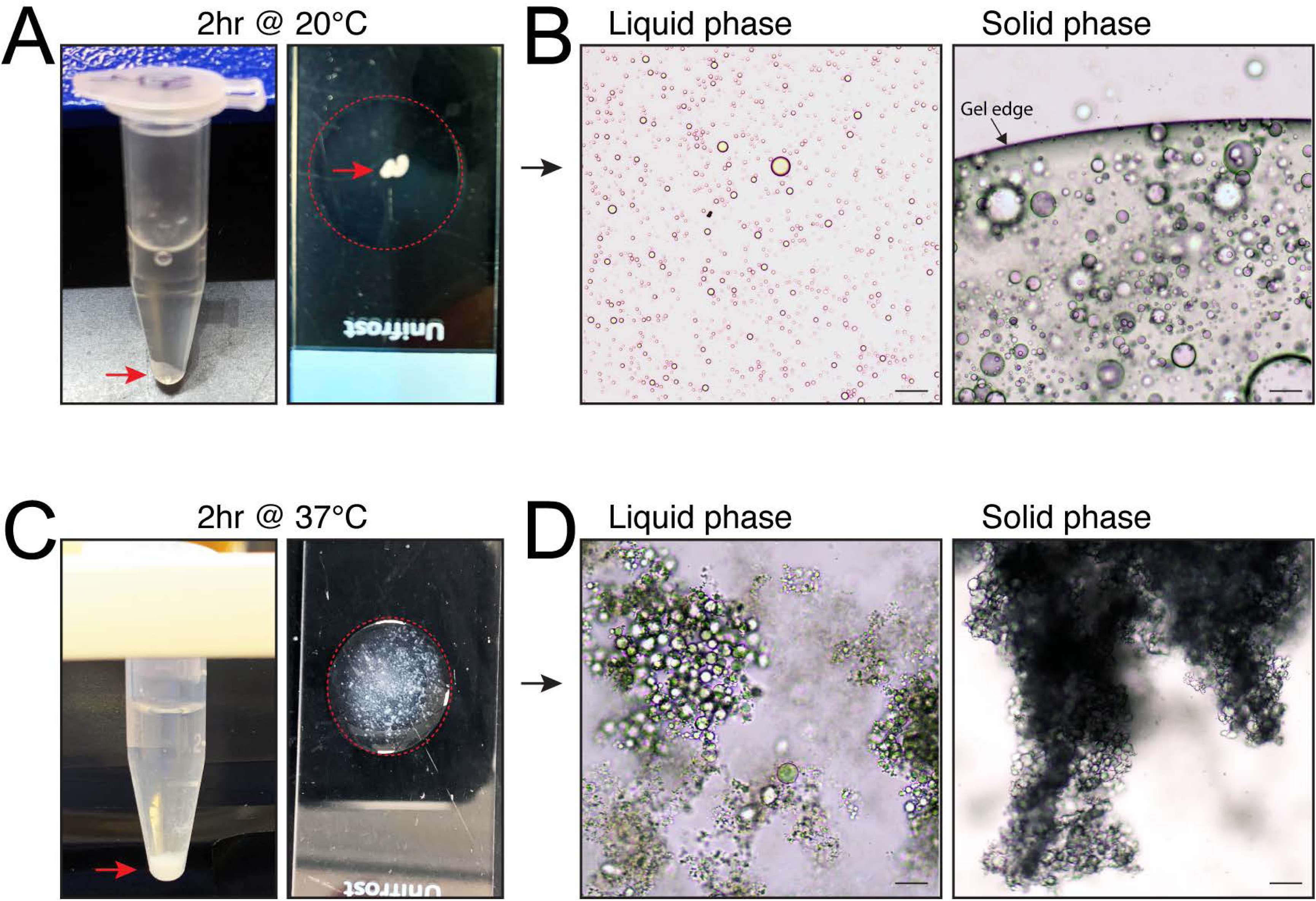
Mud condensates undergo liquid-solid transition. **(A)** Images of Mud^1955^ condensate formation reaction (100μM protein in 100mM NaCl, 20mM Tris pH 8) after 2hr incubation at 20°C showing formation of a semi-translucent solid (*left*, *red arrow*). Removal of this solid via pipetting with a cut tip allowed transfer to a microscope slide (*right*, *red arrow*). Note that the solid is soaked in surrounding solution from the reaction tube (*dashed red circle*). **(B)** *Left:* Microscopic imaging of the bulk reaction solution from (A, *left*) showing that the liquid phase (i.e. non-solid) contains numerous spherical droplets that do not undergo readily apparent fusion events. *Right:* Microscopic imaging of the gel-like solid shows it contains densely coalesced droplets of variable size. These droplets are immobile and resistant to fusions despite their close proximity. The interface between the solid and surrounding solution (‘Gel edge’ with *black arrow*) shows a sharp, distinct boundary. Scale bars, 10μm. **(C)** Images of Mud^1955^ condensate formation reaction (100μM protein in 100mM NaCl, 20mM Tris pH 8) after 2hr incubation at 37°C showing formation of a flocculent, white solid (*left*, *red arrow*). Removal of this solid via pipetting with a cut tip allowed transfer to a microscope slide (*right*, *red arrow*). Note that this dispersed solid is soaked in surrounding solution from the reaction tube (*dashed red circle*). **(D)** *Left:* Microscopic imaging of the bulk reaction solution from (A, *left*) showing that the liquid phase (i.e. non-solid) contains mostly spherical droplets that appear in clumps. *Right:* Microscopic imaging of the flocculent, white solid shows it contains more densely clumped droplets that form with an irregular border. Scale bars, 10μm.

To determine if Mud^1955^ solid formation is temperature-sensitive, we next examined reaction behavior at 37°C. We again observed formation of a solid; however, at this higher temperature the solid had a white, flocculent appearance and was easily dispersed by gentle pipetting (Figure 2C). Despite its gross resemblance to precipitated protein, microscopic examination of this solid surprisingly showed its makeup to be that of numerous phase-separated droplets clumped together in large aggregates (Figure 2D). Individual droplets within these aggregates formed more irregular spherical morphologies. Unlike the gel-like solid above, which had a sharp, defined edge, these white solids had irregular borders defined by the extent of droplet clumping. Additional imaging revealed apparent congealing of droplets within the central core of the aggregates (Figure S2B). The surrounding solution of these reactions also showed extensive phase separation, although with droplets maintaining a more spherical shape and forming smaller aggregates (Figure 2D). Overall, we conclude that Mud^1955^ condensates undergo liquid-solid transitions.

### Structural modeling implicates interdomain self-interactions in Mud that regulate condensate behavior

Having identified the coalescence and liquid-solid transition behaviors of Mud^1955^ condensates, we next sought to define how these might be regulated. Our previous biochemical studies identified a direct interaction between the Mud^CC^ and Mud^PBD^ domains, which is prevented by phosphorylation of S1868 that is within a Mud^CC^ consensus Warts kinase motif^8^. We first used AlphaFold modeling to identify the structural determinants of this interaction, using an input of three Mud^1955^ sequences to model a trimeric assembly our previous work suggested^8,25^. In agreement with these studies, this model shows a direct interaction between Mud^CC^ and the Mud^PBD^ peptide, which forms a small α-helix that packs against the coiled-coil core (Figure 3A; note that one trimeric coiled-coil binds all three Pins-binding domain peptides at three equivalent sites). This interdomain interaction is accommodated by the unstructured, flexible ‘Linker’ sequence (Mud^LINKER^) between the two domains. Notably, the predicted site of Mud^PBD^ binding within the Mud^CC^ lies adjacent to the S1868 phosphorylation site (Figure 3A).

**Figure 3.**
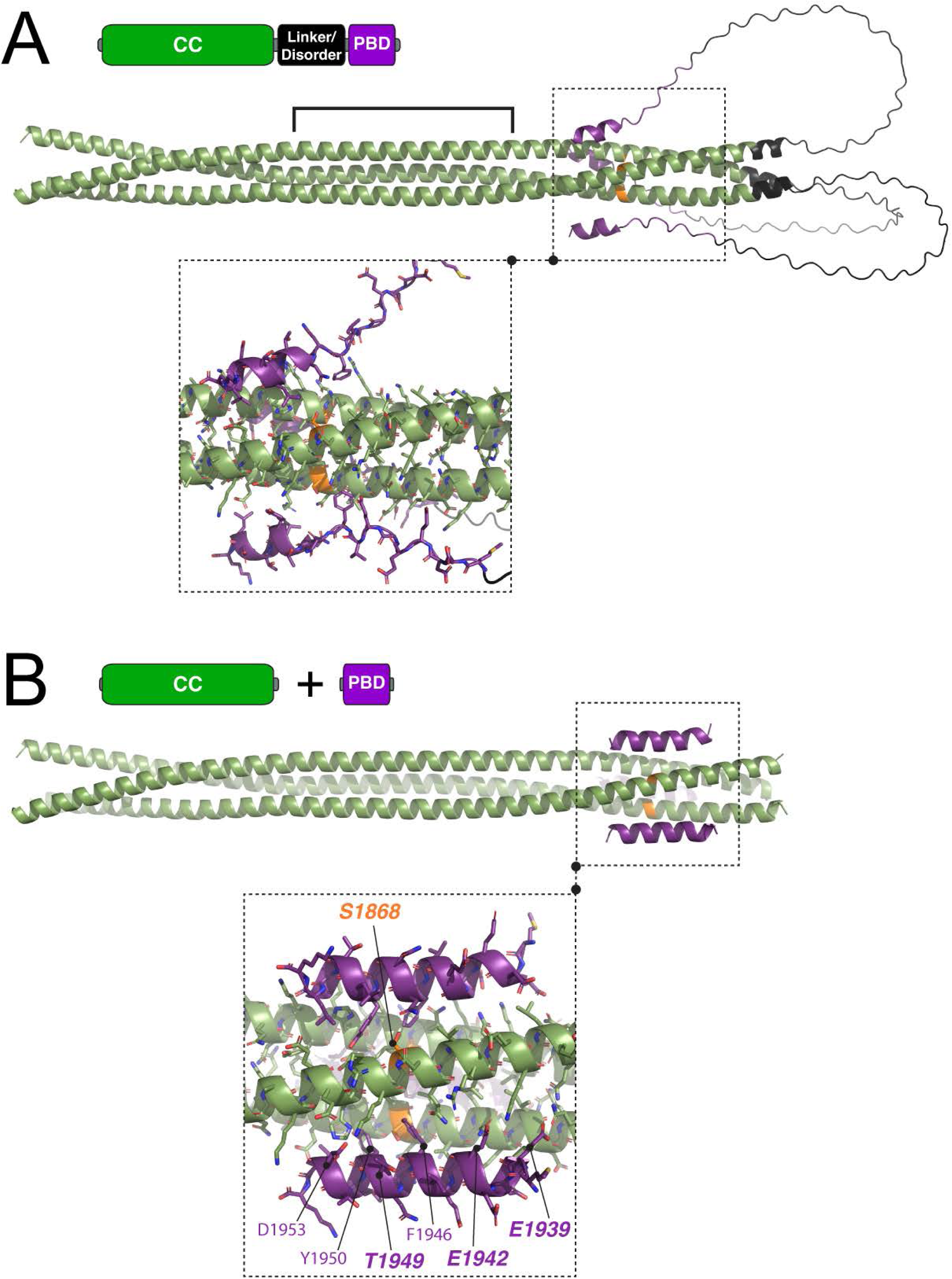
AlphaFold modeling shows interdomain Mud self-interaction. **(A)** Trimeric input of the Mud^1955^ sequence in AlphaFold produced a model with pTM=0.55, which may be negatively impacted by an overall non-idealized CC packing and assembly^60^. Specifically, the parallel CC trimer (*green*) showed idealized heptad packing in two residue segments, 1778-1809 and 1849-1881, leaving a central region of residues 1810-1848 in a non-canonical assembly (*solid bracket*). The three Mud^LINKER^ segments (*black*) each adopted an unstructured loop, which was positioned backward allowing *in cis* interaction of each Mud^PBD^ peptide (*purple*) with three identical binding sites within the idealized packing of the C-terminal region of Mud^CC^. This interaction site contained residue S1868 (*orange*) that we previously found as a substrate for Warts kinase^8^. Below is shown a zoom view of the interaction between Mud^CC^ and Mud^PBD^ segments (*dashed box*). **(B)** Trimeric input of the Mud^CC^ sequence (residues 1760-1890) together with separate input of the Mud^PBD^ sequence (residues 1938-1955) in AlphaFold produced a model with pTM=0.62. The Mud^CC^ (*green*) assumed an identical parallel trimeric assembly as in (A), again with the idealized heptad packing interrupted within the central core. Despite the lack of tethering Mud^LINKER^ segments, each of the three Mud^PBD^ domains (*purple*) showed *in trans* interaction with a nearly identical Mud^CC^ binding site as in (A). Zoomed view (*dashed box*) again demonstrates the proximity of Mud^PBD^ binding to S1868 (*orange*). Mud^PBD^ contacting residues are labeled, including E1939, E1942, and T1949 studied further herein.

We next performed additional AlphaFold modeling using the minimal Mud^CC^ and Mud^PBD^ sequences as separate inputs (lacking the Mud^LINKER^ sequence) to produce a model of the complex *in trans*. This analysis showed a similar inter-domain complex as the *in cis* model, also finding that the Mud^PBD^ binds as an α-helix packed against the S1868 motif within Mud^CC^ (Figure 3B; note that again a single trimeric coiled-coil domain binds three Pins-binding domain peptides at three equivalent sites). This interaction further corroborates the binding site identified within the Mud^CC^ domain in both structural models. Notable Mud^PBD^ residues at the binding interface include E1939 and E1942, which our previous biochemical studies found were critical for interaction^8^, along with T1949 that is within a putative Polo kinase phosphorylation motif (Figure 3B and see more below). Lastly, modeling of the corresponding sequences in NuMA, the functional mammalian Mud homolog, also showed direct interaction between the NuMA^CC^ and NuMA^LGN^ (LGN-binding domain) regions (Figure S3). A majority of NuMA^LGN^ residues in contact with NuMA^CC^ in this model are known to be involved in LGN binding^30^, suggesting a possible conserved mechanism with Mud.

To assess the involvement of S1868 phosphorylation in condensate behavior, we examined phase separation of a S1868D phosphomimetic Mud^1955^. This construct readily formed condensate droplets, indicating phosphorylation does not prevent Mud phase separation. However, in contrast to wild-type, the S1868D protein did not display the ‘droplet-of-droplets’ coalescence behavior, nor did it undergo liquid-solid phase transitioning over time (Figure 4C and data not shown). Rather, S1868D showed a higher propensity for liquid-like droplet fusions compared to wild-type (Figure 4C). Notably, AlphaFold modeling of a Mud^S1868D^ trimer found a complete disruption of the interdomain interaction seen in wild-type (Figure 4B). Thus, the direct binding between Mud^CC^ and Mud^PBD^ appears to be necessary for droplet coalescence and solid formation, with S1868 phosphorylation acting as a negative regulator of these phenotypes. We also examined an E1939A/E1942A double mutant in Mud^PBD^, which we previously showed prevents Mud^CC^ binding^8^, and the structural modeling conducted here further indicated are located at the binding interface (Figure 3B). Although this mutant protein produced condensates, as with S1868D, the droplets showed extensive liquid-like fusions and failed to undergo a solid transition compared to wild-type (Figure 5B). Together, these results suggest that the interdomain interactions of Mud^CC^ and Mud^PBD^ are critical for specific condensate behavior and that Warts kinase may be an important regulator.

**Figure 4.**
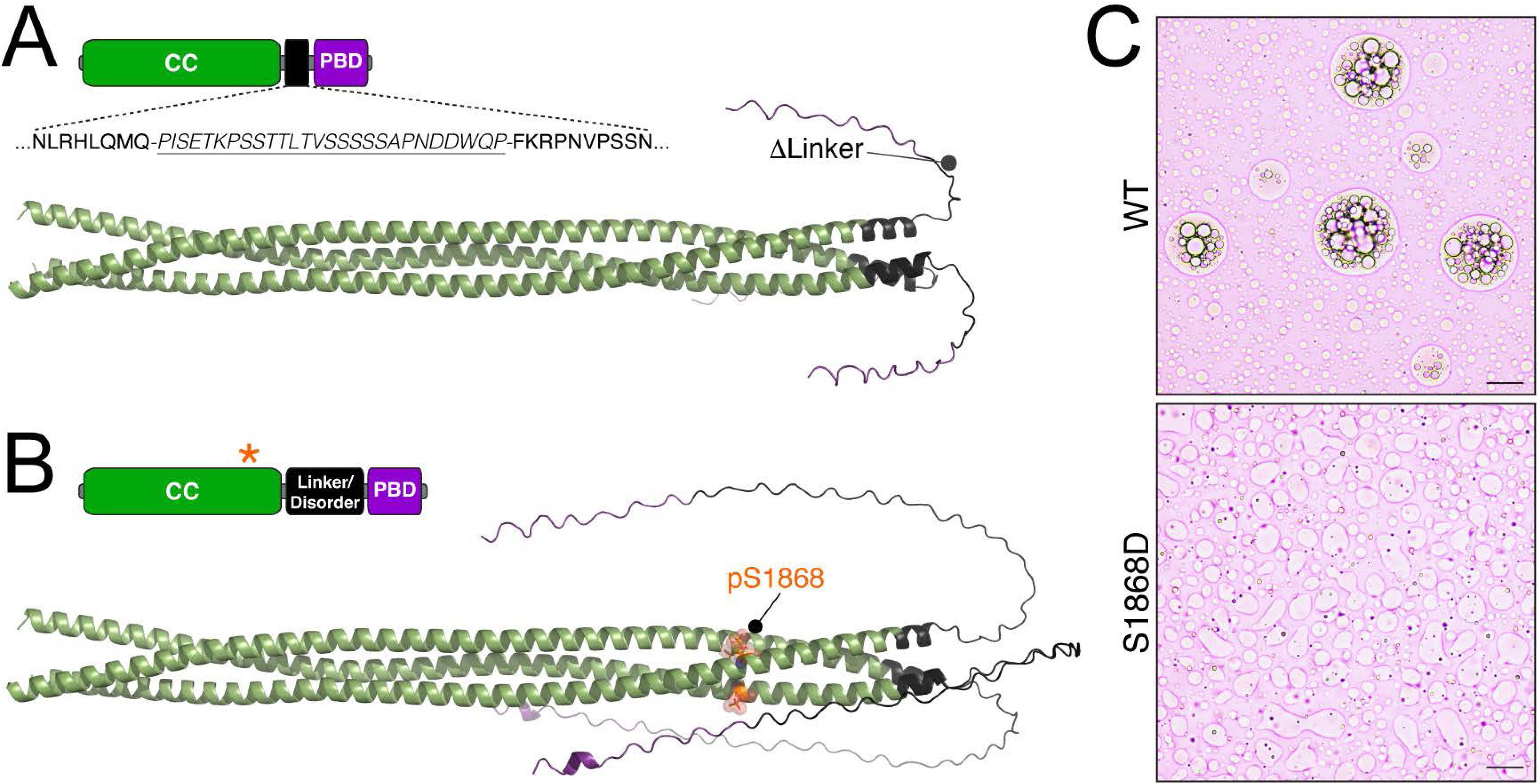
Mud^LINKER^ and S1868 are important sequence determinants for interdomain interaction and condensate behavior. **(A)** Trimeric input of the Mud^1955^ sequence, with residues 1899-1925 removed (shown in underlined and italicized text), in AlphaFold produced a model with pTM=0.58. The Mud^CC^ (*green*) assembled similarly to Mud^1955^, but the truncated Mud^LINKER^ sequences (*black*) does not permit interaction of the Mud^PBD^ (*purple*) with Mud^CC^. **(B)** Trimeric input of the Mud^1955^ sequence, with S1868 marked for modeling of phosphorylation (pS1868, *orange*), in AlphaFold produced a model with pTM=0.57. The Mud^CC^ (*green*) again assembled similarly to Mud^1955^; however, phosphorylation of Mud^CC^ residue S1868 within the predicted binding site for Mud^PBD^ (*purple*) uncouples the interdomain interaction. No alterative interaction between either Mud^PBD^ or Mud^LINKER^ with Mud^CC^ is predicted in this model. **(C)** Images of Mud^1955^ condensate formation reactions (100μM protein in 100mM NaCl, 20mM Tris pH 8) for wild-type protein (WT; *top*) and the phosphomimetic mutant S1868D (*bottom*). The wild-type protein forms stable spherical condensates that do not undergo extensive fusion events. Also seen are several ‘droplet of droplets’ assemblies formed from apparent droplet coalescence. In contrast, the S1868D protein produces condensates that readily undergo dynamic liquid fusions as evident from the distorted droplet shapes. Scale bars, 10μm.

**Figure 5.**
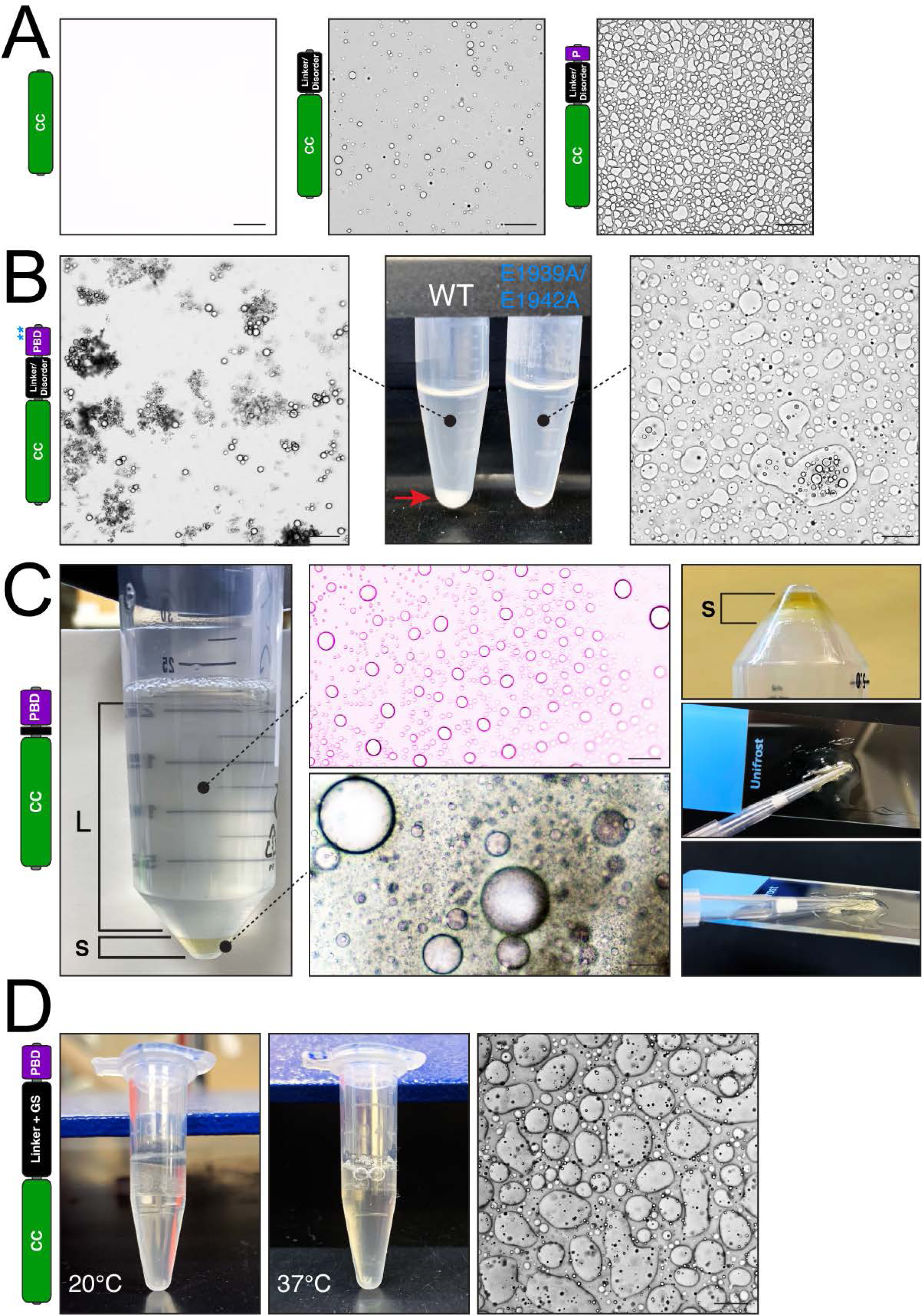
Mud^LINKER^ and interdomain association influence behavior of Mud condensates. **(A)** *Left:* The isolated Mud^CC^ protein does not produce condensates. *Middle:* A protein that includes the Mud^CC^ and Mud^LINKER^ produces small, spherical droplets. *Right:* Mud protein including residues 1760-1947 produces dynamic condensates that undergo liquid-like fusions evident by their distorted shapes. All reactions were conducted under identical conditions (100μM protein in 100mM NaCl, 20mM Tris pH 8; at 20°C). Scale bars, 10μm. **(B)** Condensate formation reactions of Mud^1955^ proteins (100μM protein in 100mM NaCl, 20mM Tris pH 8) were conducted at 37°C. *Left:* Microscopic image of soluble fraction of Mud^1955^ wild-type protein reaction shows clusters of small spherical droplets. *Middle:* Gross images of condensate formation reactions. Whereas wild-type (WT) protein produces a white solid (*red arrow*), the double mutant E1939A/E1942A (*blue text* and see *blue \*\** in cartoon indicating their relative positions) does not. *Right:* Microscopic image of E1939A/E1942A mutant protein reaction shows formation of dynamic condensates. Scale bars, 10μm. **(C)** *Left:* Gross image of Mud^ΔLINKER^ during purification (prior to final concentration steps; contained in 300mM NaCl, 20mM Tris pH 8). Spontaneous phase separation occurred overnight at 4°C, resulting in a visible gel-like solid (S) and the remaining bulk liquid (L) phase. *Middle-top:* Microscopic image of the liquid phase shows formation of spherical condensates. *Middle-bottom:* Microscopic image of the solid phase shows coalescence of spherical condensates of varying sizes. *Right-top:* Gross image of the bulk solid phase following removal of liquid phase and exposure to air drying. *Right-middle* and *-bottom:* Gross images of small amounts of solid phase following air drying demonstrates a hardening effect capable of gluing pipette tip to microscope slide. Note that in both images the slide is suspended by holding the pipette tip (not shown) rather than resting on a surface. Scale bars, 10μm. **(D)** *Left:* Gross image of Mud^LINKER+GS^ condensate formation reaction at 20°C demonstrating the lack of solid formation. *Middle:* Similarly, gross image of Mud^LINKER+GS^ condensate formation reaction at 37°C also demonstrating the lack of solid formation. *Righ:* Microscopic image of Mud^LINKER+GS^ condensate formation reaction at 20°C demonstrating the formation of liquid-like droplets with distorted shapes indicative of dynamic fusion events. Scale bars, 10μm.

### Critical role for LINKER-PBD sequences in Mud condensate behavior

We next investigated the role of the Mud^LINKER^ region in condensate formation and behavior. This was motivated by the structural modeling suggesting this unstructured, flexible motif permits the intramolecular Mud complex formation (Figure 3A), as well as the nature of this sequence itself, which contains a repetitive stretch of small polar amino acids (particularly serine) known to participate in phase separation of other proteins^31,32^. We first examined Mud^CC^ alone, lacking both the Mud^LINKER^ and Mud^PBD^ sequences, which failed to produce any condensates (Figure 5A). Addition of the Mud^LINKER^ sequence resulted in the formation of small, spherical condensate droplets. Further extension to include the N-terminal segment of the Mud^PBD^ sequence that was not predicted to make direct contacts with Mud^CC^ resulted in more extensive condensate formation, which showed a highly dynamic liquid-like property (Figure 5A). Notably, neither of these constructs produced the coalesced or solid phenotypes seen with the full Mud^1955^ sequence, although they nevertheless indicate a key role for the Mud^LINKER^ in condensate formation.

As noted above, a conspicuous sequence tract of polar residues lies within Mud^LINKER^ (Figure 4A). As similar sequences have been implicated in solid transitions in other biological condensates^31,32^, we next examined how deletion of this segment of Mud^LINKER^ within Mud^1955^, while retaining the full Mud^PBD^ sequence (termed Mud^ΔLINKER^ herein), impacts condensate function. Notably, AlphaFold modeling of this construct showed a loss of intramolecular association between Mud^CC^ and Mud^PBD^ domains, presumably due to a loss of flexible sequence space required to adopt complex formation (Figure 4A). Surprisingly, Mud^ΔLINKER^ spontaneously underwent extensive phase separation during the purification process, prior to final concentration and while still at 300mM NaCl ionic strength (note wild-type does not form condensates at this NaCl concentration, see Figure 1B). Moreover, a liquid-solid transition was evident during purification as well (Figure 5C). While the bulk liquid solution contained isolated spherical condensate droplets, the gel-like solid showed coalesced spherical droplets of varying size (Figure 5C). This phenotype was similar to that seen in the full Mud^1955^ protein, although the Mud^ΔLINKER^ solid contained more extensive coalescence, did not require reduction in NaCl concentration to form (300mM compared to 100mM), and presumably occurred at a lower protein concentration (the spontaneous solid formation prior to final purification steps prevented an accurate measurement). The Mud^ΔLINKER^ solid also had a denser consistency, and while conducting imaging experiments of it we found that samples allowed to dry produced a hardened, glue-like solid (Figure 5C). These findings indicate that this polar sequence span within Mud^LINKER^ (residues 1899-1925) is not strictly required for condensate formation or solid transition. Based on the structural modeling of Mud^ΔLINKER^, they also suggest that intramolecular Mud^CC^/Mud^PBD^ domain interactions are also likely to be dispensable for these properties.

To further address these implications, we also examined how increasing the flexibility and sequence space of Mud^LINKER^ impacts condensate activity by inserting a glycine-serine (GS) repeat (x12) between Mud^LINKER^ and Mud^PBD^ regions in an otherwise native Mud^1955^ background. Unlike its Mud^ΔLINKER^ counterpart, this Mud^LINKER+GS^ protein did not form a gel-like solid either during purification or upon further examination at conditions promoting Mud^1955^ solid formation (Figure 5D). It did, however, produce condensates that showed liquid-like fusions over time (Figure 5D). Thus, truncation of the interdomain linker sequence enhances solid transition of Mud condensates, whereas extending its length and flexibility prevents this phenomenon. As mentioned above, the E1939A/E1942A mutant that uncouples Mud domain self-association also leads to a loss of solid transition and produces condensates with more dynamic liquidity (Figure 5B). Lastly, we tested the sufficiency of the Mud^LINKER^ sequence to induce condensate formation by fusing it to GFP. This GFP^LINKER^ protein did not show condensate formation and appeared indistinguishable from GFP alone control. Thus, while Mud^LINKER^ appears to play a role in Mud condensate formation and behavior, it is not sufficient to induce these effects. Altogether, these results suggest that the interdomain interaction between Mud^CC^ and Mud^PBD^ plays a critical role in the liquid-solid condensate transition, with the connecting Mud^LINKER^ sequence providing an important accessory role in this process.

### Increasing multivalency of Mud self-interactions impacts condensate formation

The striking phenotype seen with Mud^ΔLINKER^ (Figure 5C-D), together with structural modeling indicating that this construct fails to form an intramolecular complex (Figure 4A), raised the possibility of instead an intermolecular interaction wherein the Mud^PBD^ of one Mud^1955^ molecule might associate with the Mud^CC^ of a second. Extrapolation of this model (i.e. the second Mud molecule binding a third, and so forth) could implicate Mud homooligomerization as a possible molecular explanation for condensate behavior. To test this model further, we designed a Mud construct that appended two additional LINKER-PBD sequences to the Mud^1955^ protein (termed Mud^PBDx3^ herein). Formation of an intramolecular complex should only involve a single PBD sequence binding *in cis* at a given Mud^CC^ binding site, as shown in Figure 3. In the event of intermolecular assembly, each of the multiple PBD sequences could bind *in trans* to a separate Mud^CC^, increasing the multivalency of the resultant Mud complexes. We found that Mud^PBDx3^ formed phase separated condensates when examined at protein concentration 10- or 100-fold below the standard concentration used for Mud^1955^, which did not phase separate under these lower concentration conditions (Figure 6). Somewhat surprisingly, however, Mud^PBDx3^ did not form the coalesced droplets seen with Mud^1955^ at higher concentration, instead displaying a highly dynamic, liquid-like behavior (Figure 6). This may be due to increased flexibility in the assembled complexes provided by the additional Mud^LINKER^ sequences. Despite this distinction, these results are consistent with the valency of Mud complexes contributing to condensate formation, which agrees with other phase separating proteins described previously^33^, and supports a model of Mud assembly involving intermolecular oligomerization following *in trans* interactions between Mud^CC^ and Mud^PBD^ domains.

**Figure 6.**
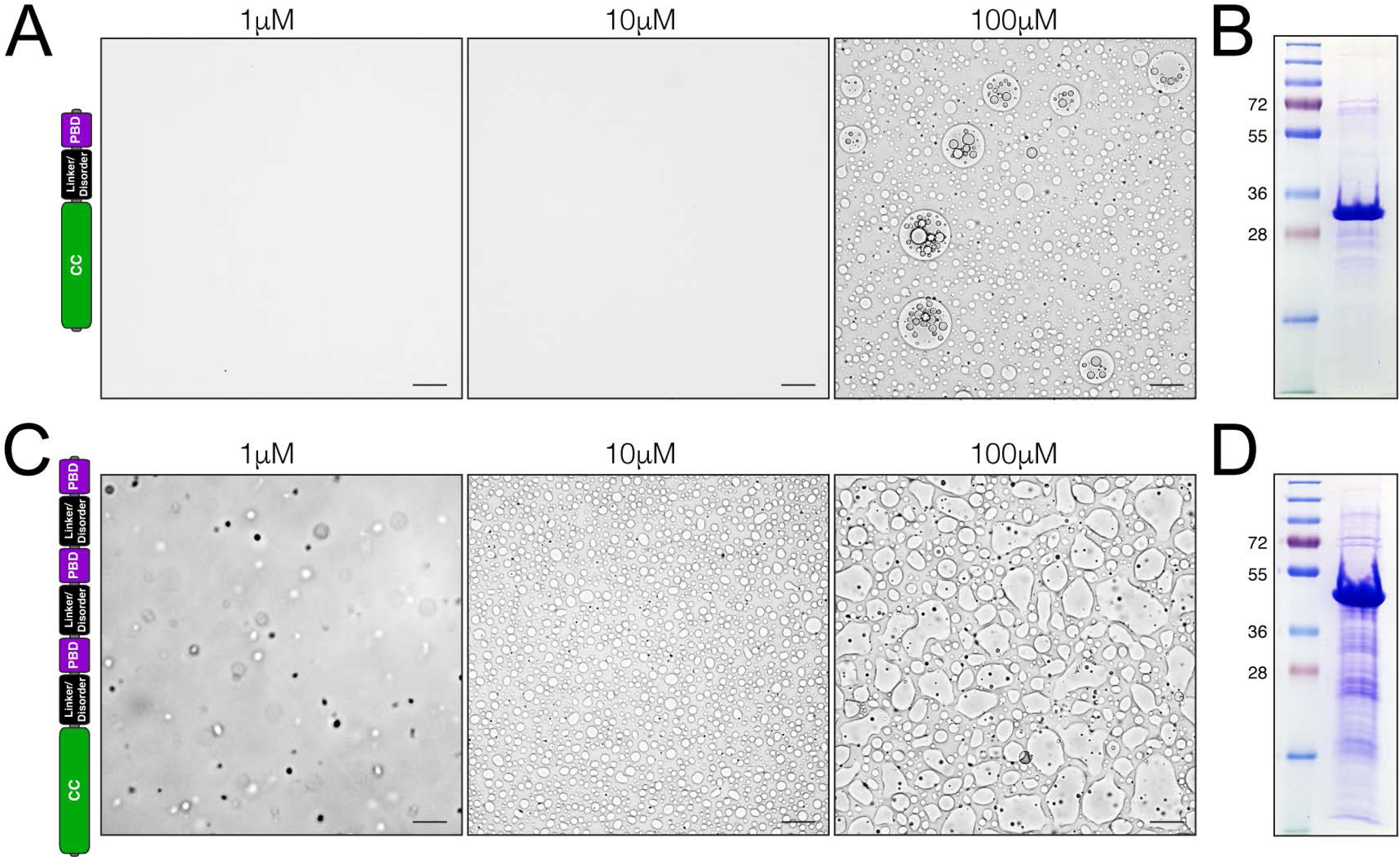
Increasing multivalency potential of interdomain associations enhances Mud condensate formation. **(A)** Indicated concentrations of Mud^1955^ (100mM NaCl, 20mM Tris pH 8) were imaged for condensate formation and morphology. At 100μM, an abundance of stable, spherical condensates are present, along with several coalesced ‘droplet of droplets’. At both lower concentrations (1μM and 10μM), no condensates are apparent. Scale bars, 10μm. **(B)** SDS-PAGE analysis of Mud^1955^ protein used in (A). **(C)** Indicated concentrations of Mud^PBDx3^ (100mM NaCl, 20mM Tris pH 8) were imaged for condensate formation and morphology. At 1μM, small spherical droplets are visible. At 10μM, larger condensates form and show evidence of droplet fusions. At 100μM, condensates form that undergo extensive droplet fusions. Scale bars, 10μm (except 1 μM protein, where Scale bar, 1μm). **(D)** SDS-PAGE analysis of Mud^PBDx3^ protein used in (C).

### Plk1 phosphorylates Mud^PBD^ and regulates condensate dynamics

Having identified sequence determinants that impact condensate activity of Mud^1955^, we explored additional mechanisms that could regulate their properties. With Mud^CC^ S1868 phosphorylation already having shown to increase condensate liquidity (Figure 4), we sought to identify additional phosphorylation sites focusing here on Mud^PBD^. Using the Eukaryotic Linear Motif (ELM) server for *in silico* sequence analysis (http://elm.eu.org/), we identified a putative Polo-like kinase 1 (Plk1; Polo in flies) site within the C-terminal end of the Mud^PBD^ sequence (T1949; Figure 7A). Radiometric ^32P^ATP assays using purified Plk1 kinase and Mud^1955^ substrate showed robust phosphorylation signal (Figure 7B). Mutation of T1949 to a non-phosphorylatable glutamate (T1949E) resulted in a total loss of phosphorylation, indicating that this is the sole Plk1 site within Mud^1955^.

**Figure 7.**
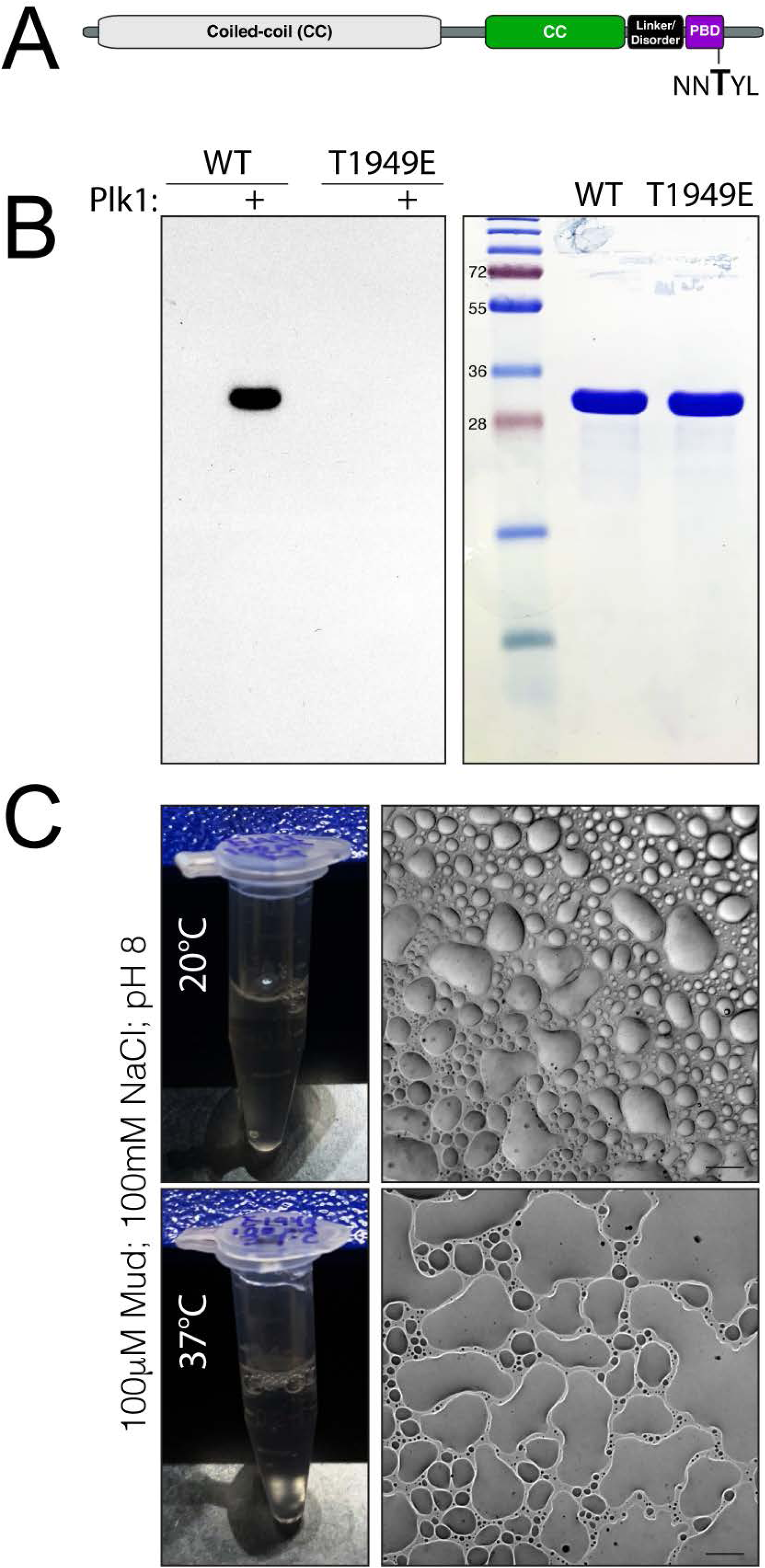
Plk1 phosphorylates Mud^PBD^ and increases liquidity of Mud condensates. **(A)** Domain architecture of Mud indicating the Plk1 consensus motif within Mud^PBD^. Phosphorylation is predicted to occur at residue T1949 (*bold*). **(B)** *Left:* Mud^1955^ wild-type (WT) or T1949E mutant (20μg) were incubated in the absence or presence (+) of 1μg of Plk1 for 30 minutes at 30°C. Shown is the autoradiographic detection of ^32^P phosphate incorporation indicating a strong phosphorylation of WT that is absent in T1949E. *Right:* Coomassie stained SDS-PAGE gel shows equivalent loading amounts and purity of the Mud proteins used. **(C)** *Top-left:* Gross image of Mud^T1949E^ condensate formation reaction with indicated conditions following 2hr incubation at 20°C shows no solid formation. *Top-right:* Microscopic image shows Mud^T1949E^ condensates have a liquid-like behavior. *Bottom-left* and *-right:* Identical imaging but instead from condensate formation reactions at 37°C. No solid transition is apparent, and condensates formed show more prominent liquidity and surface wetting. Scale bars, 10μm.

We then examined the phase separation properties of the T1949E phosphomimetic mutant (Mud^T1949E^). This protein showed extensive droplet formation; however, unlike the wild-type protein, Mud^T1949E^ readily underwent liquid-like fusions over time (Figure 7C). Mud^T1949E^ did not show any coalesced droplet phenotype, nor did it undergo any evident solid transition over time at either 20°C or 37°C (Figure 7C). Overall, these data identify a Plk1 phosphorylation site within Mud^PBD^ and suggest this modification could play a critical role in regulating the biophysical properties of Mud condensates.

### Additional coiled-coil spindle-associated proteins produce solid condensates in vitro

Recent studies identified a subset of spindle-associated proteins existing within a highly dynamic, liquid-like phase separated domain that facilitates assembly of the meiotic spindle in mammalian oocytes^34^. Similar to Mud, many of these and other spindle-associated proteins contain coiled-coil domains, which are known to facilitate phase separation in diverse proteins^35^. Thus, we sought to compare with Mud the phase separation behavior of CC domain-containing proteins identified in this liquid-like meiotic spindle domain. For the most direct Mud comparison, we chose two centrosomal proteins for these initial analyses, the Transforming Acidic Coiled-Coil protein (TACC) and NudE (Figure 8).

**Figure 8.**
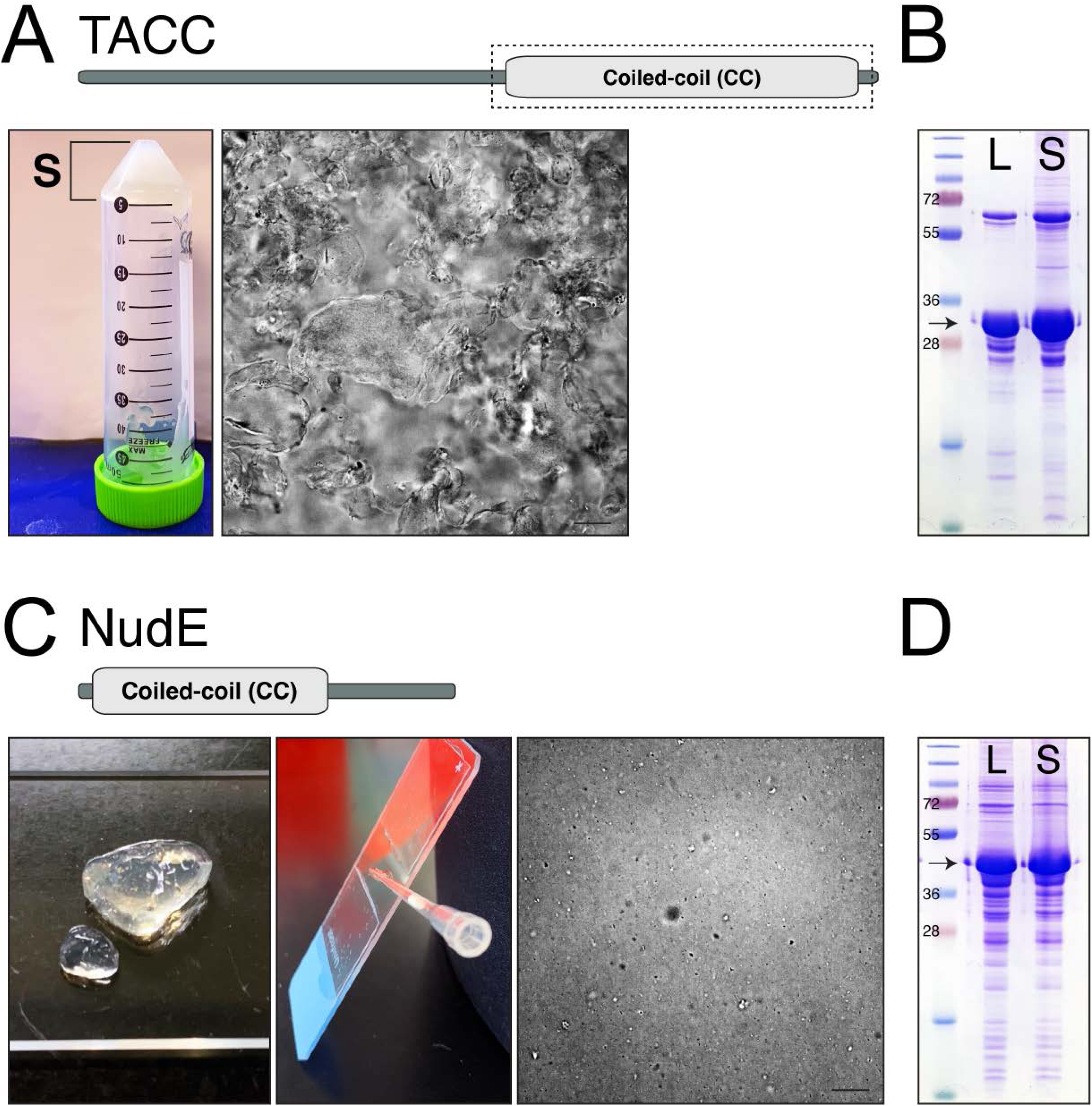
Additional spindle pole coiled-coil proteins, TACC and NudE, form solids. **(A)** *Left:* Purification of the coiled-coil (CC; *top*) domain from TACC results in formation of a solid (S; note that the liquid phase has been removed in the image shown). *Right:* Microscopic imaging of the TACC solid shows a dense, amorphous network lacking any discernable condensate droplets as seen in Mud solids. Scale bars, 10μm. **(B)** SDS-PAGE analysis of TACC protein used in (A). The solid (S) sample shows similar protein identity as the remaining liquid (L) phase. **(C)** *Left:* Purification of full-length NudE (*top*) results in formation of a solid (S) with gross properties consistent with a hydrogel. *Middle:* Drying of the NudE solid forms a hardened solid capable of gluing a pipette tip to the microscope slide. *Right:* Microscopic imaging of the NudE solid shows a dense structure lacking any discernable condensate droplets as seen in Mud solids. Scale bars, 10μm. **(D)** SDS-PAGE analysis of NudE protein used in (C). The solid (S) sample shows similar protein identity as the remaining liquid (L) phase.

Attempts to purify a full-length TACC protein were hampered by solubility and purity issues, which prompted us to examine a C-terminal region containing the CC domain. Similar to Mud^ΔLINKER^, this TACC^CC^ spontaneously separated into a solid-like material during its purification (Figure 8A), which was revealed to contain the TACC^CC^ protein similar to the soluble fraction that remained (Figure 8B). Imaging of the TACC^CC^ solid, however, did not reveal coalesced droplets as seen with Mud, but rather a dense, compact material (Figure 8A). In contrast to TACC, expression and purification of a full-length NudE was possible. Similar to TACC^CC^, NudE underwent a spontaneous phase separation during purification, producing a gel-like solid that remained intact with physical manipulation and contained NudE protein similar to the remaining soluble phase (Figure 8C-D). Imaging of this solid also did not reveal the presence of coalesced droplets as with Mud^1955^; however, when allowed to dry the NudE solid became a stiff, glue-like solid similar to Mud^ΔLINKER^ (Figure 8C). These properties, along with the gross inspection of the NudE solid, are consistent with protein-based hydrogels previously described for both naturally occurring and *de novo* designed sequences^36–38^. These results suggest that coiled-coil domain proteins functioning at spindle poles may have a common property of solid-forming phase separation.

## DISCUSSION

Mud is a critical regulator of Dynein activity, controlling mitotic spindle assembly and orientation essential for proper cell division. While the spindle orientation function of Mud at the cell cortex occurs in collaboration with Pins in certain cell types (e.g. neuroblasts), this activity in epithelial cells is Pins-independent as is Mud function at spindle poles across cell types^6,12,22^. Delineating how these Pins-dependent and -independent Mud functions are differentially controlled is of particular relevance to understanding its distinct mitotic roles. Our results highlight a potential role for unique aspects of phase separated biological condensates, here defining the key sequence determinants for the formation and behavior of homotypic Mud condensates, which differ from our previous studies of Mud/Pins complex condensates^25^. Several notable conclusions can be drawn: (1) neither Mud^CC^ nor Mud^LINKER-PBD^ is sufficient for condensate formation, (2) addition of the Mud^LINKER^ sequence to Mud^CC^ produces liquid-like condensates, (3) further extension to include the Mud^PBD^ (e.g. Mud^1955^) produces condensates with reduced droplet fusions that instead coalesce and undergo liquid-solid transition, and (4) manipulation of the Mud^LINKER^ sequence alters the liquid-solid dynamics of condensates with a shortened sequence yielding more significant gelation and an extended sequence producing more liquid-like droplets that fail to form solids. The coalesced ‘droplet-of-droplets’, along with the solid transition, are distinctive traits of homotypic Mud condensates that sharply contrast with the highly dynamic, liquid-like behavior of Mud/Pins condensates^25^. We propose that these biophysical differences in condensates may contribute to the distinct functions of the cortical Mud/Pins complex and the Pins-independent Mud activity at spindle poles.

Liquid-solid condensate phase transition appears to be driven by direct, self-interactions between Mud^CC^ and Mud^PBD^ domains, supported by their mutual requirement and the loss of solid formation in the E1939A/E1942A and phosphomimetic mutants that are each predicted to decouple this interaction (Figures 4, 5, and 7 and see^8^). More specifically, we envision that intermolecular interaction among multiple Mud molecules acts as a molecular driver of condensate behavior. Assembly of the coiled-coil domain necessarily generates a Mud molecule with multiple PBD sequences (modeled herein as three from trimeric assembly), presumably each capable of binding another identically assembled Mud molecule. Such associations would lead to formation of a multivalent Mud oligomer that may underlie the behavior of the condensates produced. Artificially increasing the multivalency potential through additional PBD sequences did not enhance solid transition, but it did facilitate condensate formation at lower concentrations (Figure 6). Previous studies in other systems have spotlighted a role for multivalent interactions in condensate formation and phase transitions^39,40^. Liquid-solid transitions also typically involve intrinsically disordered regions, such as in the FUS protein, and are often associated with pathological outcomes^41,42^. The Mud^LINKER^ region has an abundance of polar amino acids consistent with protein disorder, and its fusion with Mud^CC^ was sufficient to induce condensate formation in the absence of Mud^PBD^ (Figures 4 and S4). However, deletion of this sequence exacerbated solid formation indicating it is not required for this property of Mud condensates. Taken together, these results suggest the Mud^LINKER^ sequence itself contributes to condensate formation, but that its length, by determining the distance and flexibility between the Mud^CC^ and Mud^PBD^ domains, dictates condensate dynamics.

We further discovered that solid transition of Mud condensates is negatively regulated at two phosphorylation sites recognized by the mitotic kinases, Warts and Polo (Figures 4 and 7, respectively). Notably, both of these kinases have established functions at spindle poles and are important regulators of spindle assembly and orientation^8,43–46^. Recent studies have found similar effects of phosphorylation on NuMA condensates. In this case, the mitotic kinase Aurora A phosphorylates the condensate-forming C-terminal fragment of NuMA, resulting in increased liquid-like fluidity and reduced solid-like transitions^28,47^. Aurora A activity is cell cycle dependent, with activation prominent in mitosis where it facilitates spindle assembly and function^48^. Its activity is proposed to increase the liquid properties of NuMA condensates in mitotic cells, which may be critical for localization from spindle poles to adjacent microtubules and the cell cortex^28^. Whether Mud is phosphorylated by Aurora A is unknown, but *aurA* mutants have altered cortical Mud and spindle orientation defects despite normal spindle pole localization in neuroblasts^49,50^. Our results suggest that Warts or Polo may provide a similar regulatory function regarding the dynamics of Mud condensates.

Finally, put in context with our previous studies, another regulatory aspect of Mud condensates appears to be its association with the cortical adaptor protein Pins. Whereas homotypic Mud condensates show limited liquid-like dynamics and form solids over time, those of Mud/Pins complexes undergo extensive droplet fusions leading to surface wetting over time and do not undergo solid transitions^25^. Notably, previous studies on NuMA-dependent microtubule bundling, critical for spindle assembly and pole focusing, found this activity to be inhibited by its binding to LGN (the mammalian Pins homolog)^13^. Moreover, the same NuMA C-terminal region governing its phase separation performs this microtubule bunding activity, and this overlaps with the Mud sequence used in our studies herein. Thus, although specific details are likely to differ, it is apparent that similar dynamics and regulatory mechanisms control Mud and NuMA phase separation. We propose that the Mud^PBD^ plays a crucial role in determining condensate activity– when bound to Pins, dynamic liquid-like Mud/Pins condensates are produced; when self-associating with additional Mud molecules, coalesced condensates result that are capable of solid formation. These distinctions may ultimately produce differential Mud activity that influence Dynein-dependent and -independent function^5,28,51–55^.

### LIMITATIONS OF THE STUDY

A notable limitation of our study is its use of a relatively small Mud sequence fragment rather than an extended sequence or, ideally, a full-length protein. We attempted studies with larger Mud fragments, including with the extended coiled-coil region N-terminal to the Mud^CC^ examined here (Figure 1A). Unfortunately, these were hampered by poor expression and solubility of these constructs. However, it should be noted again that the Mud^1955^ sequence used in this study is coded by exons uniquely included in the *mud* isoform that regulates spindle positioning and other microtubule activities^12,29^. Another limitation of our study is its focus on Mud function *in vitro*. Additional studies exploring Mud condensate formation and behavior *in vivo* will be important future directions. Additional future directions could include further biophysical characterization of Mud solids, as well as expanding studies on TACC and NudE along with additional coiled-coil proteins known to regulate spindle function. Lastly, NudE is known to participate in Mud/Pins-mediated spindle orientation through interactions with 14-3-3 protein complexes^56^, and examining if this interaction alters NudE phase separation remains another important open question.

## STAR METHODS

## Key resources table

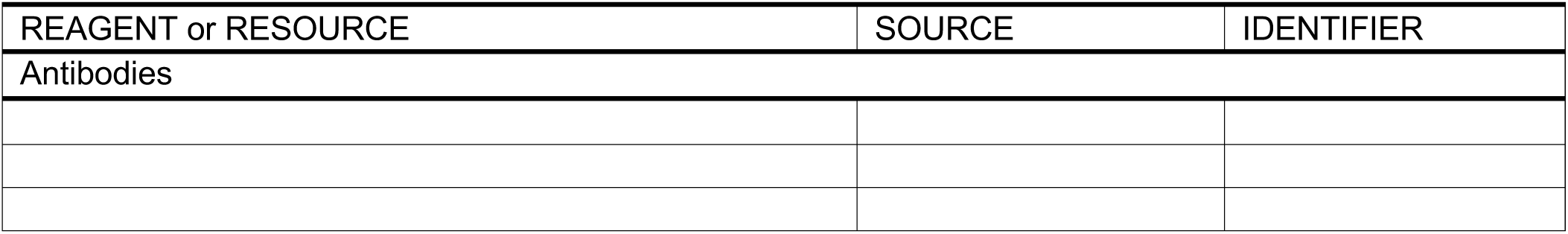
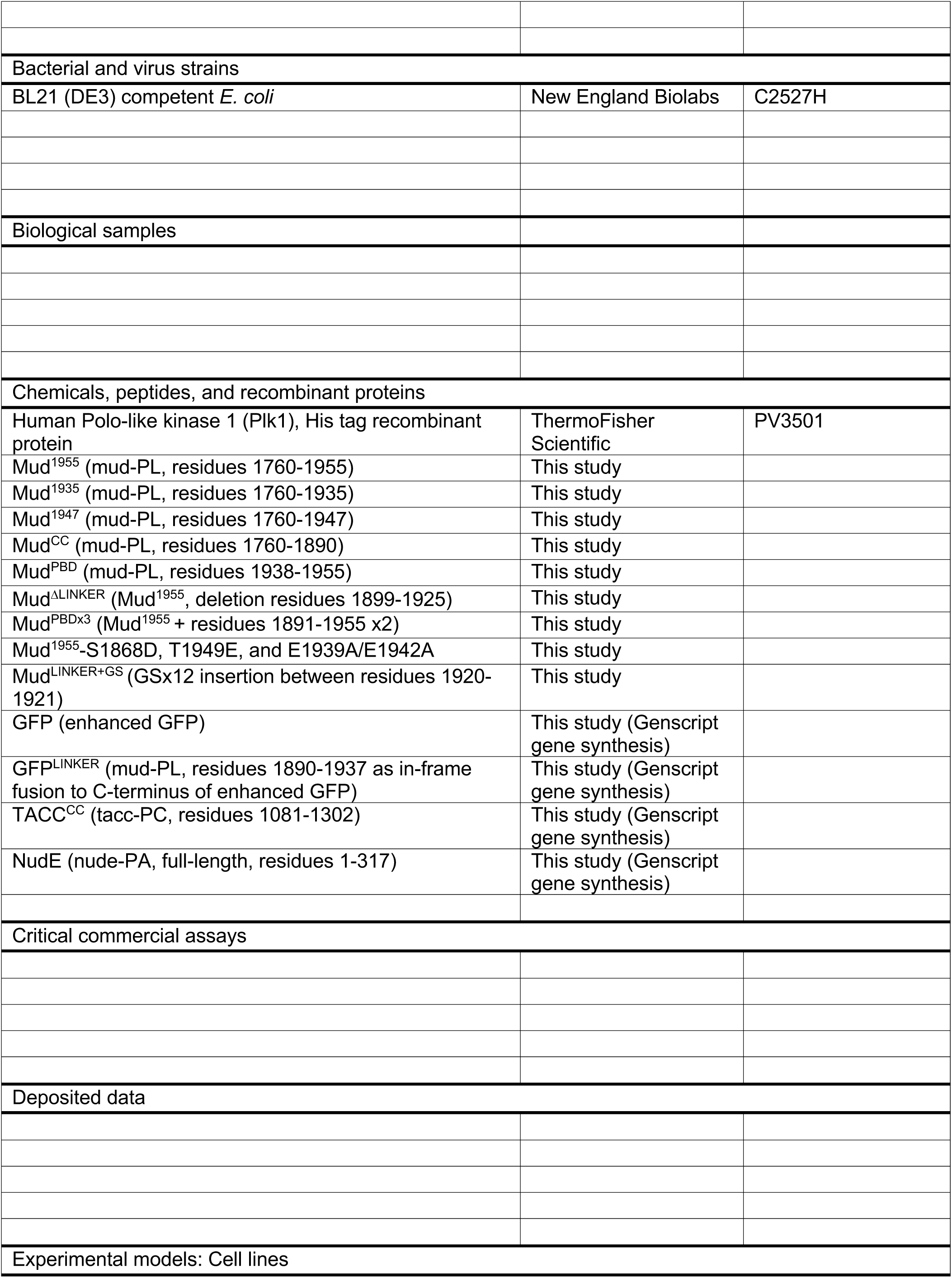
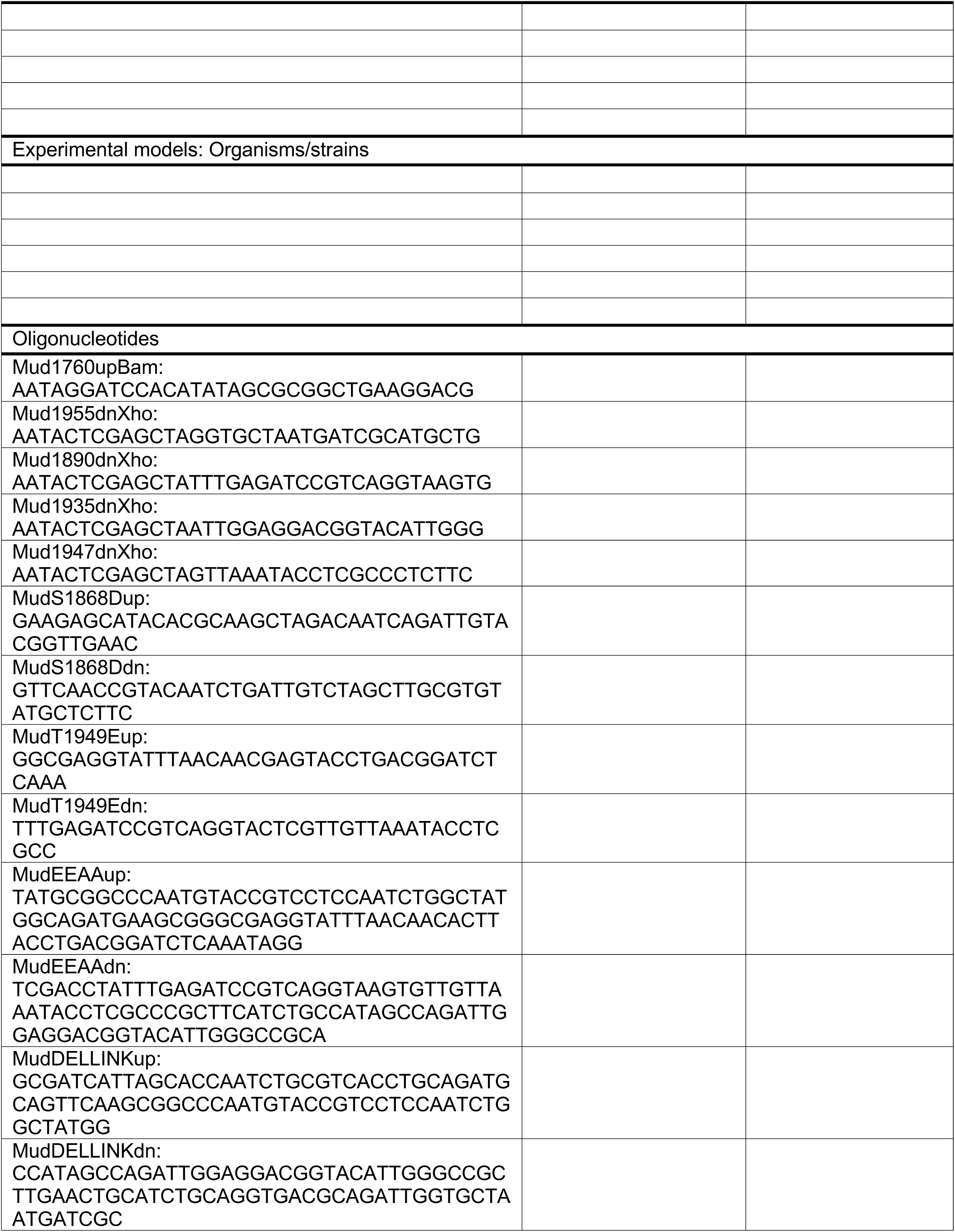
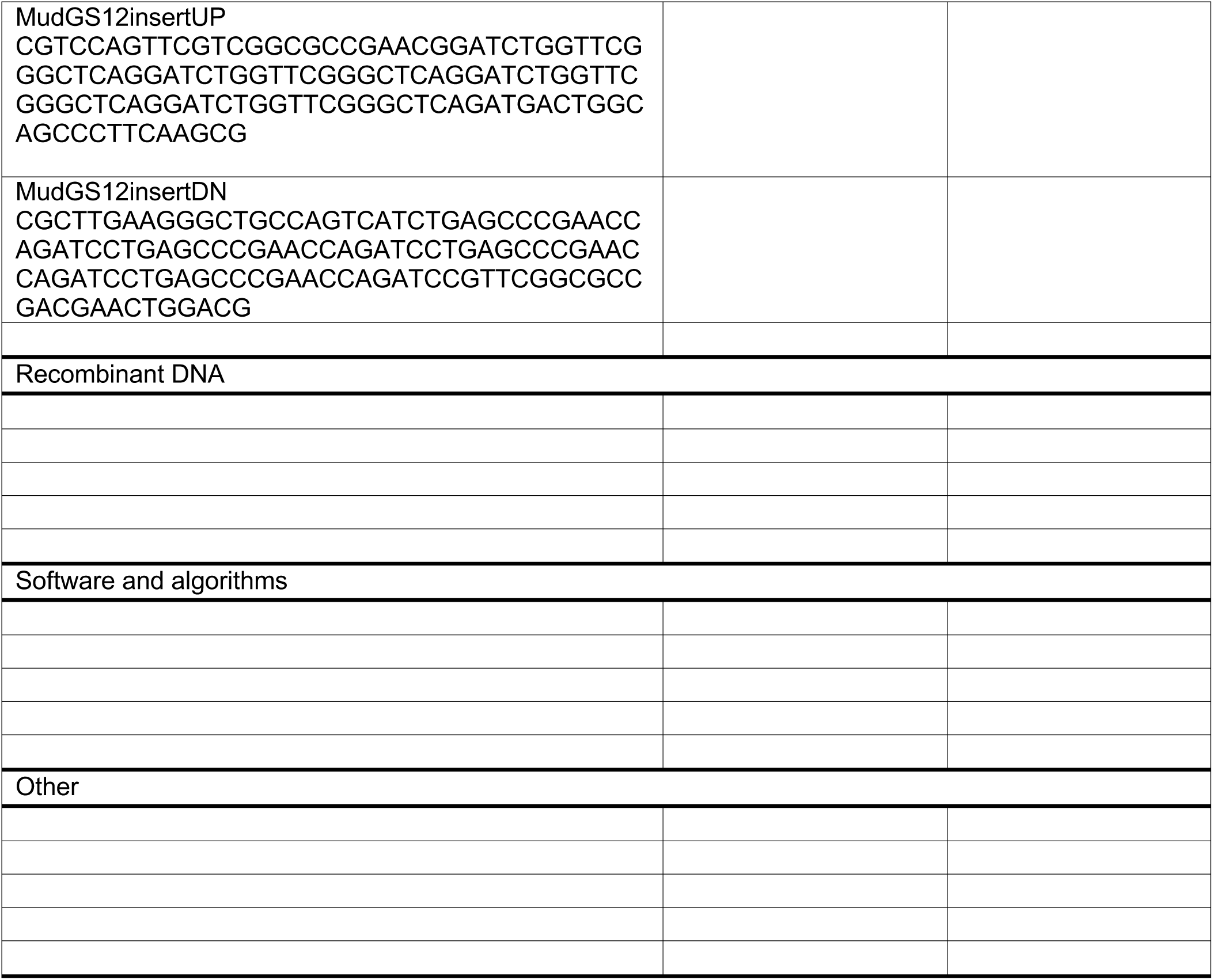

## Resource availability

Further information and requests for resources should be directed to and will be fulfilled by the lead contact, Dr. Christopher Johnston

## Materials availability

Requests for bacterial expression plasmids should be directed to Dr. Christopher Johnston.

## Method details

### Molecular cloning

cDNA cloning for bacterial protein expression was performed using PCR amplified fragments obtained from a *Drosophila* S2 cell cDNA library template and cloned into the pBH plasmid with 5′-*BamH*I or -*Bgl*II and 3′-*Xho*I or *Sal*I restriction sites. Site-directed, deletional, and insertional mutageneses were carried out with a standard PCR protocols using KOD-XL DNA polymerase (EMD Millipore, catalog #71087). The sequence was generated in the pUC57 cloning vector by Genewiz (Azenta Life Sciences, South Plainfield, NJ, USA) and subsequently subcloned into pBH as described above.

### Protein purification

All proteins were expressed in BL21(DE3) *E. coli* under induction of isopropyl β-D-1-thiogalactopyranoside (IPTG) and grown in standard Luria–Bertani broth supplemented with 100 μg/ml ampicillin. Transformed cells were grown at 37°C to an OD_600_ ∼0.6 and induced with 0.2 mM IPTG overnight at 20°C. Cells were harvested by centrifugation (5000 × *g* for 10 min), and bacterial pellets were resuspended in lysis buffer and flash-frozen in liquid nitrogen. Cells were lysed using a Branson digital sonifier and clarified by centrifugation (12,000 × g for 30 min). Lysis was performed in N1 buffer (50 mM Tris pH8, 500 mM NaCl, 10 mM imidazole) and coupled to Ni-NTA resin (Qiagen, catalog #30210) for 3 h at 4°C. Following extensive washing in N1, proteins were eluted with N2 buffer (50 mM Tris pH8, 500 mM NaCl, 300 mM imidazole). The 6×His tag was removed using TEV protease during overnight dialysis into N1 buffer. Cleaved products were reverse affinity purified by a second incubation with Ni-NTA resin and collection of the unbound fraction. Final purification was carried out using an S200-sephadex size exclusion column equilibrated in storage buffer (20 mM Tris pH8, 200 mM NaCl, 2 mM DTT).

### Phase separation assays

Proteins were diluted at indicated concentrations in minimal buffer without addition of molecular crowding agents (20mM Tris pH 7.5, 100mM NaCl; unless otherwise noted) and incubated at indicated temperatures and for indicated times (typically room temperature for 5 minutes unless otherwise noted). Subsequently, 150 μL of reactions were added to glass microscope slides in chambers assembled from affixed coverslip spacers. Coverslips were applied and samples were imaged to visualize the formation of LLPS droplets using an Olympus CX43 brightfield microscope with an Olympus EP50 camera for imaging. All images shown are representative of at least 3 independent experiments.

### Kinase phosphorylation assays

Human Polo-like kinase 1 (Plk1), His tag recombinant protein was purchased from ThermoFisher Scientific (Waltham, MA, USA). Mud^1955^ (10 μg; wild-type or T1949E), with or without Plk1 (1 μg), was diluted in ice-cold assay buffer (20 mM Tris [pH 7.4], 100 mM NaCl, 1 mM DTT, 10 mM MgCl_2_, and 10 μM ATP). ATP-γ-^32^P (5 μCi) was added at 30°C for 30 min. Reactions were quenched by addition of SDS loading buffer. Samples were resolved by SDS-PAGE and dried on filter paper using a BioRad 583 gel dryer. Phosphorylation, determined by ^32^P incorporation, was assessed using autoradiographic film processing (Kodak Biomax). Images shown are representative of at least 3 independent experiments.

### AlphaFold 3 structural modeling

Protein structural modeling was performed using the AlphaFold server (www.alphafoldserver.com) utilizing the current AlphaFold 3 code^57^. Primary protein sequences were input along with the desired copy number (e.g. ‘3’ for prediction and modeling of trimeric coiled-coil assemblies). Confidence metrics were interpreted for model accuracy. Structural models were assessed using per-atom confidence (pLDDT) and predicted template modeling (pTM) scores, which assesses the accuracy of the entire structure^58,59^. All models presented scored above the pTM threshold for possible similarity to the true structure (see figure legends for specific scorings). Models were further interpreted and shown images were rendered in PyMOL (www.pymol.org).

## FIGURE LEGENDS

**Figure S1.**
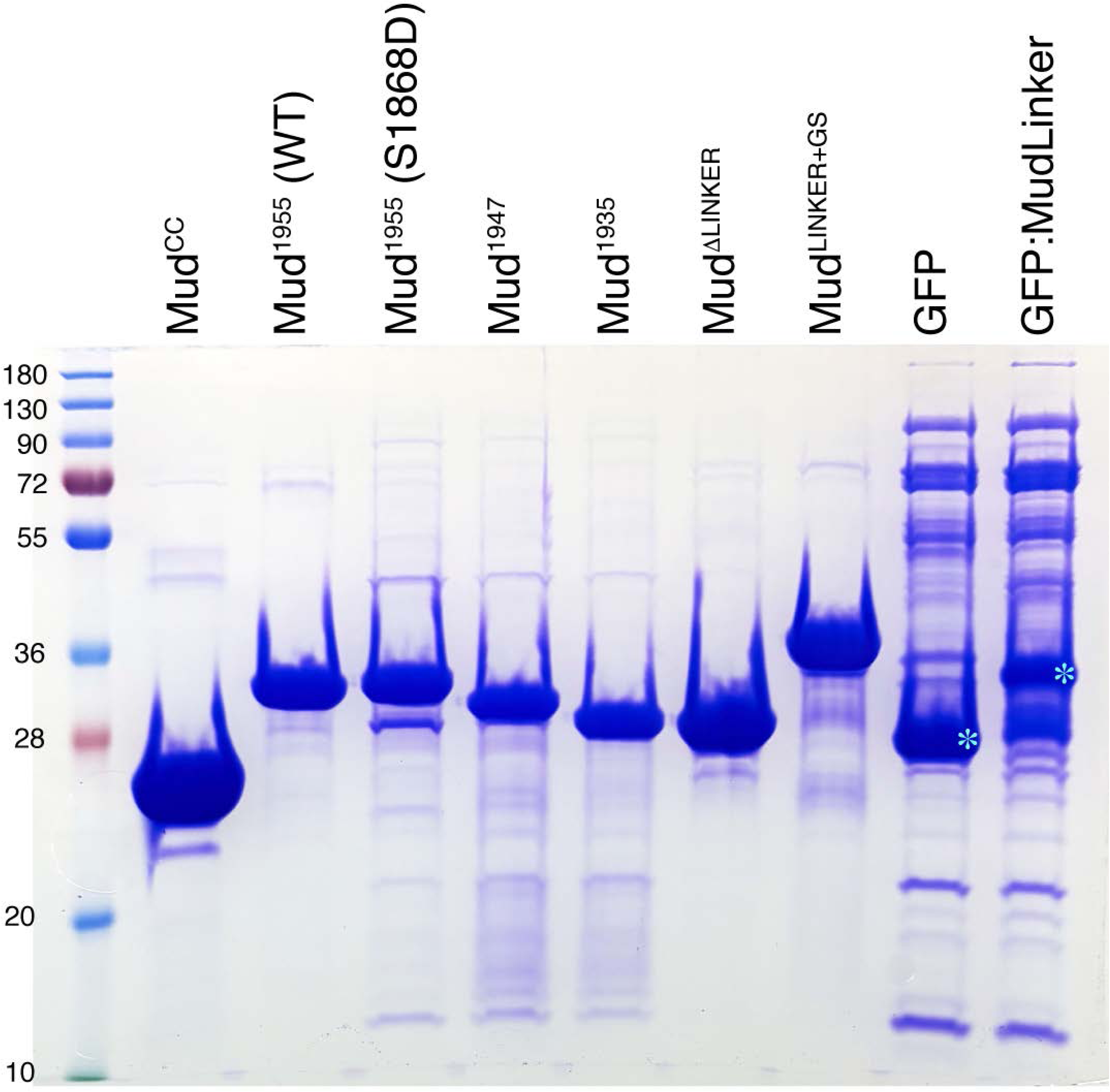
Additional spindle pole coiled-coil proteins, TACC and NudE, form solids. SDS-PAGE analysis of most proteins used in this study. Approximately 30μg of total protein was loaded in each respective lane. See individual figures in the main text for analysis of additional proteins.

**Figure S2.**
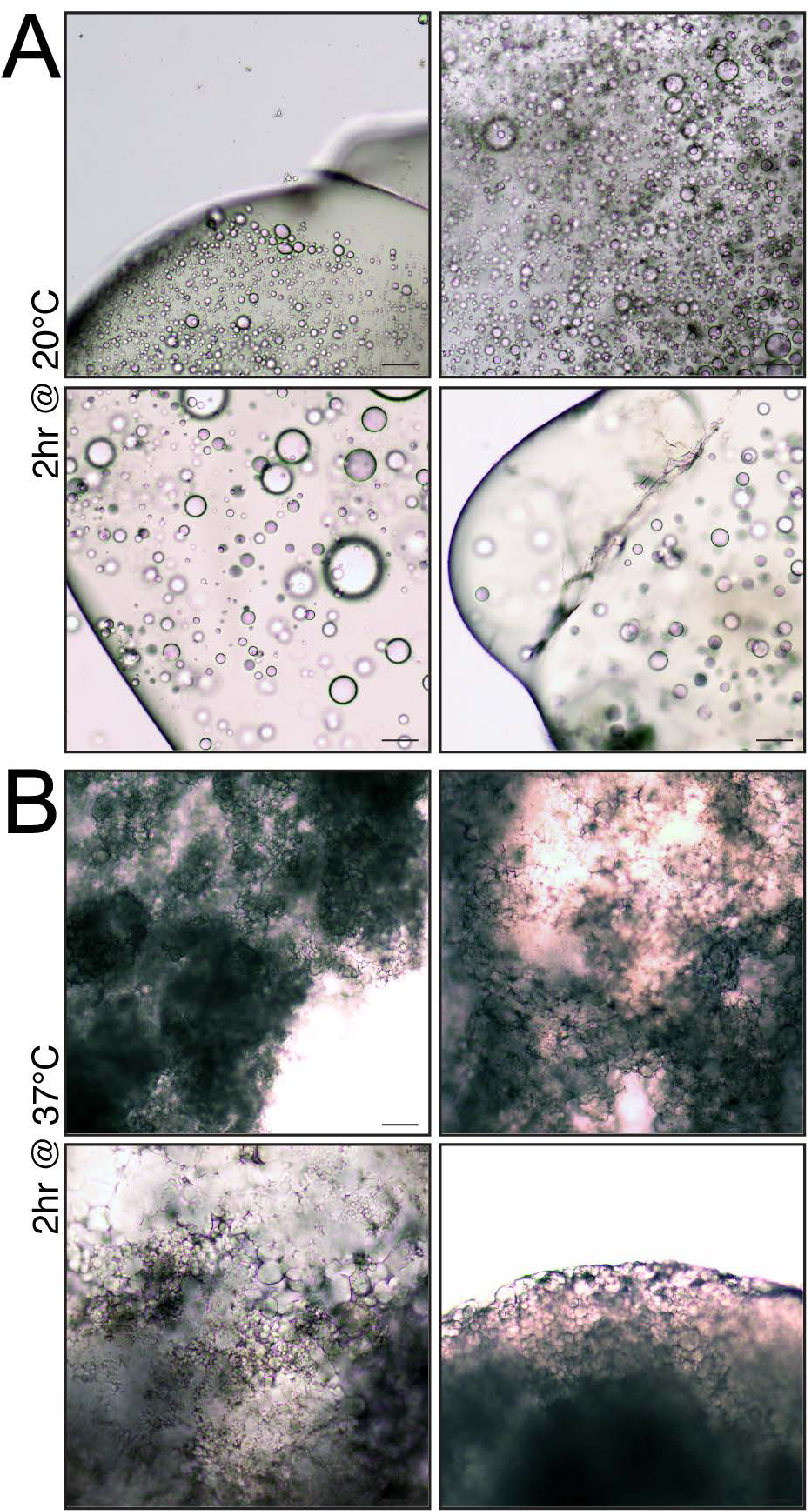
Additional imaging of Mud^1955^ solid transitions. **(A)** Shown are four additional examples of the gel-like Mud^1955^ solid formed in a condensate reaction (100μM protein in 100mM NaCl, 20mM Tris pH 8) incubated for 2hr at 20°C. The image in the *upper-right* shows entirely the interior of a solid, whereas the other three images have represented the gel edges. In all cases, the solid contains numerous spherical droplets that do not undergo liquid-like fusions. Scale bars, 10μm. **(B)** Shown are four additional examples of the flocculent white Mud^1955^ solid formed in a condensate reaction (100μM protein in 100mM NaCl, 20mM Tris pH 8) incubated for 2hr at 37°C. Scale bars, 10μm.

**Figure S3.**
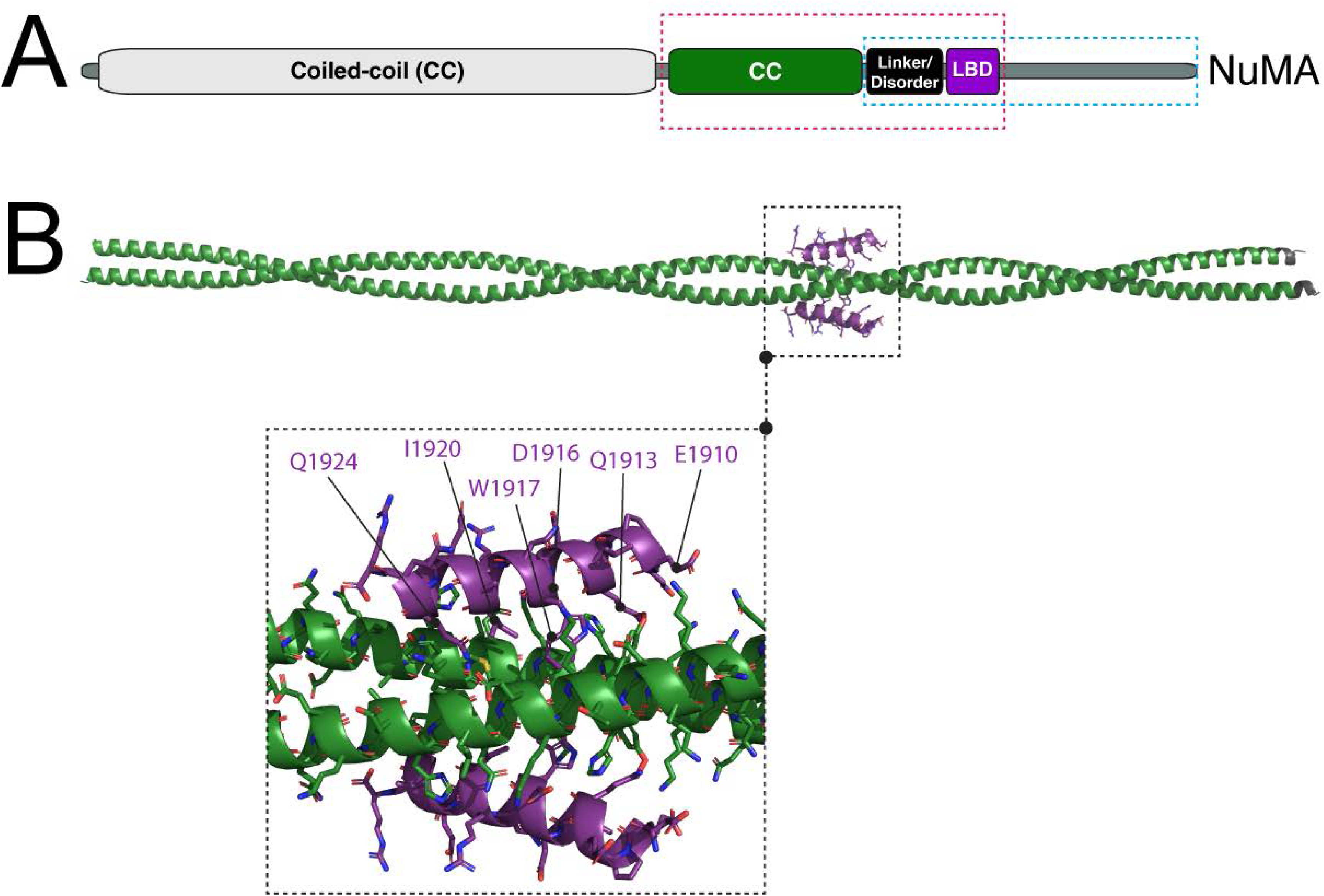
Structural modeling of a dimeric Mud1955 and the homologous sequence from human NuMA reveals conserved interdomain interactions. **(A)** Domain architecture of NuMA shows homology with Mud (see Figure 1A), having an extended N-terminal coiled-coil (CC; *white*) followed by a short CC (*green*) that precedes the LGN-binding domain (LBD; *purple*). These two domains are separated by a linker sequence of predicted disorder (Linker/Disorder; *black*). The region of NuMA homologous to Mud^1955^ is depicted with *dashed red box*. The NuMA^CC^ domain (*green*) is longer than Mud^CC^ (234 versus 130 residues) and forms a dimer^5^. The NuMA^LINKER^ region is also much longer than Mud^LINKER^ (95 versus 48 residues). The C-terminal region previously implicated in NuMA phase separation includes the Linker/Disorder and LBD and is depicted with the *dashed cyan box*^28^. **(B)** Independent dimeric inputs of the NuMA^CC^ (*green*) and NuMA^LBD^ (*purple*) sequences in AlphaFold produced a model with pTM=0.53. Modeling of a dimeric NuMA *in cis* assembly (not shown) found a similar interdomain interaction and binding site, but the confidence of this model failed to meet the threshold likely due to the extended Linker region. A zoomed view of the interaction is shown below (*dashed box*), with specific NuMA^LBD^ residues that contact NuMA^CC^ labeled, all of which (except Q1910) also bind LGN.

**Figure S4.**
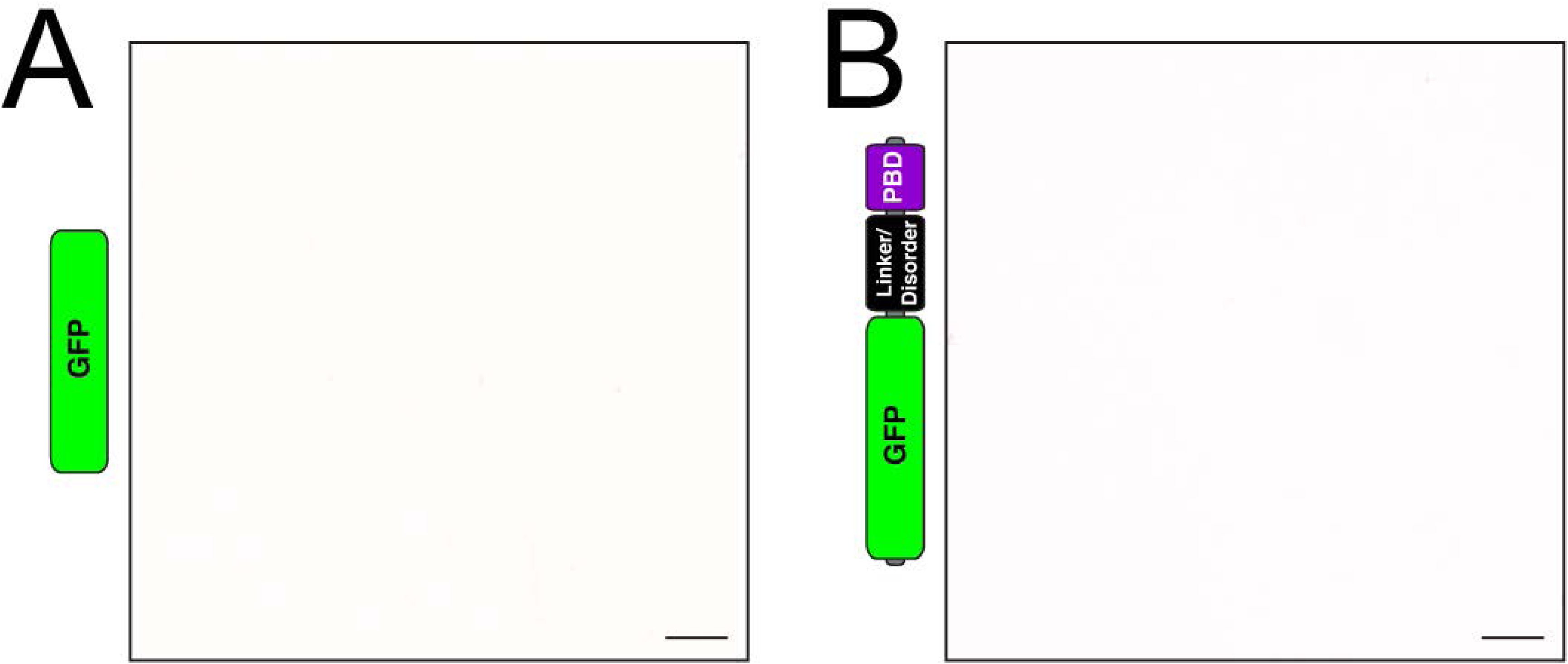
The Mud^LINKER-PBD^ is not sufficient for condensate formation. **(A)** GFP alone negative control does not form condensates (100μM protein in 100mM NaCl, 20mM Tris pH 8 at 20°C). **(B)** GFP fused to the Mud^LINKER^ and Mud^PBD^ also does not form condensates (100μM protein in 100mM NaCl, 20mM Tris pH 8 at 20°C).

## DECLARATION OF INTERESTS

The authors declare no competing interests.

## ACKNOWLEDGMENTS

Research reported in this publication was supported by the National Science Foundation (https://www.nsf.gov) under award number 2205405 (C.A.J). The content is solely the responsibility of the authors and does not necessarily represent the official views of the National Science Foundation. There was no additional external funding received for this study.

## Notes

### Competing Interest Statement

The authors have declared no competing interest.

## REFERNCES CITED

1. Valdez, V.A., Neahring, L., Petry, S., and Dumont, S. (2023). Mechanisms underlying spindle assembly and robustness. Nat Rev Mol Cell Biol 24, 523–542. 10.1038/s41580-023-00584-0.

2. Ou, G., and Scholey, J.M. (2022). Motor Cooperation During Mitosis and Ciliogenesis. Annu Rev Cell Dev Biol 38, 49–74. 10.1146/annurev-cellbio-121420-100107.

3. Kiyomitsu, T., and Cheeseman, I.M. (2013). Cortical dynein and asymmetric membrane elongation coordinately position the spindle in anaphase. Cell 154, 391–402. 10.1016/j.cell.2013.06.010.

4. Kotak, S., Busso, C., and Gönczy, P. (2012). Cortical dynein is critical for proper spindle positioning in human cells. J Cell Biol 199, 97–110. 10.1083/jcb.201203166.

5. Aslan, M., d’Amico, E.A., Cho, N.H., Taheri, A., Zhao, Y., Zhong, X., Blaauw, M., Carter, A.P., Dumont, S., and Yildiz, A. (2024). Structural and functional insights into activation and regulation of the dynein-dynactin-NuMA complex. bioRxiv. 10.1101/2024.11.26.625568.

6. Bosveld, F., Markova, O., Guirao, B., Martin, C., Wang, Z., Pierre, A., Balakireva, M., Gaugue, I., Ainslie, A., Christophorou, N., et al. (2016). Epithelial tricellular junctions act as interphase cell shape sensors to orient mitosis. Nature 530, 495–498. 10.1038/nature16970.

7. Bowman, S.K., Neumüller, R.A., Novatchkova, M., Du, Q., and Knoblich, J.A. (2006). The Drosophila NuMA Homolog Mud regulates spindle orientation in asymmetric cell division. Dev Cell 10, 731–742. 10.1016/j.devcel.2006.05.005.

8. Dewey, E.B., Sanchez, D., and Johnston, C.A. (2015). Warts phosphorylates mud to promote pins-mediated mitotic spindle orientation in Drosophila, independent of Yorkie. Curr Biol 25, 2751–2762. 10.1016/j.cub.2015.09.025.

9. Izumi, Y., Ohta, N., Hisata, K., Raabe, T., and Matsuzaki, F. (2006). Drosophila Pins-binding protein Mud regulates spindle-polarity coupling and centrosome organization. Nat Cell Biol 8, 586–593. 10.1038/ncb1409.

10. Nakajima, Y.I., Lee, Z.T., McKinney, S.A., Swanson, S.K., Florens, L., and Gibson, M.C. (2019). Junctional tumor suppressors interact with 14-3-3 proteins to control planar spindle alignment. J Cell Biol 218, 1824–1838. 10.1083/jcb.201803116.

11. Ségalen, M., Johnston, C.A., Martin, C.A., Dumortier, J.G., Prehoda, K.E., David, N.B., Doe, C.Q., and Bellaïche, Y. (2010). The Fz-Dsh planar cell polarity pathway induces oriented cell division via Mud/NuMA in Drosophila and zebrafish. Dev Cell 19, 740–752. 10.1016/j.devcel.2010.10.004.

12. Siller, K.H., Cabernard, C., and Doe, C.Q. (2006). The NuMA-related Mud protein binds Pins and regulates spindle orientation in Drosophila neuroblasts. Nat Cell Biol 8, 594–600. 10.1038/ncb1412.

13. Du, Q., Taylor, L., Compton, D.A., and Macara, I.G. (2002). LGN blocks the ability of NuMA to bind and stabilize microtubules. A mechanism for mitotic spindle assembly regulation. Curr Biol 12, 1928–1933. 10.1016/s0960-9822(02)01298-8.

14. Merdes, A., Heald, R., Samejima, K., Earnshaw, W.C., and Cleveland, D.W. (2000). Formation of spindle poles by dynein/dynactin-dependent transport of NuMA. J Cell Biol 149, 851–862. 10.1083/jcb.149.4.851.

15. Merdes, A., Ramyar, K., Vechio, J.D., and Cleveland, D.W. (1996). A complex of NuMA and cytoplasmic dynein is essential for mitotic spindle assembly. Cell 87, 447–458. 10.1016/s0092-8674(00)81365-3.

16. Silk, A.D., Holland, A.J., and Cleveland, D.W. (2009). Requirements for NuMA in maintenance and establishment of mammalian spindle poles. J Cell Biol 184, 677–690. 10.1083/jcb.200810091.

17. Bosveld, F., Ainslie, A., and Bellaïche, Y. (2017). Sequential activities of Dynein, Mud and Asp in centrosome-spindle coupling maintain centrosome number upon mitosis. J Cell Sci 130, 3557–3567. 10.1242/jcs.201350.

18. Ito, A., and Goshima, G. (2015). Microcephaly protein Asp focuses the minus ends of spindle microtubules at the pole and within the spindle. J Cell Biol 211, 999–1009. 10.1083/jcb.201507001.

19. Du, Q., Stukenberg, P.T., and Macara, I.G. (2001). A mammalian Partner of inscuteable binds NuMA and regulates mitotic spindle organization. Nat Cell Biol 3, 1069–1075. 10.1038/ncb1201-1069.

20. Johnston, C.A., Hirono, K., Prehoda, K.E., and Doe, C.Q. (2009). Identification of an Aurora-A/PinsLINKER/Dlg spindle orientation pathway using induced cell polarity in S2 cells. Cell 138, 1150–1163. 10.1016/j.cell.2009.07.041.

21. Williams, S.E., Beronja, S., Pasolli, H.A., and Fuchs, E. (2011). Asymmetric cell divisions promote Notch-dependent epidermal differentiation. Nature 470, 353–358. 10.1038/nature09793.

22. Bergstralh, D.T., Lovegrove, H.E., Kujawiak, I., Dawney, N.S., Zhu, J., Cooper, S., Zhang, R., and St Johnston, D. (2016). Pins is not required for spindle orientation in the Drosophila wing disc. Development 143, 2573–2581. 10.1242/dev.135475.

23. Kiyomitsu, T., and Boerner, S. (2021). The Nuclear Mitotic Apparatus (NuMA) Protein: A Key Player for Nuclear Formation, Spindle Assembly, and Spindle Positioning. Front Cell Dev Biol 9, 653801. 10.3389/fcell.2021.653801.

24. Radulescu, A.E., and Cleveland, D.W. (2010). NuMA after 30 years: the matrix revisited. Trends Cell Biol 20, 214–222. 10.1016/j.tcb.2010.01.003.

25. Parra, A.S., Moezzi, C.A., and Johnston, C.A. (2023). Drosophila Adducin facilitates phase separation and function of a conserved spindle orientation complex. Front Cell Dev Biol 11, 1220529. 10.3389/fcell.2023.1220529.

26. Banani, S.F., Lee, H.O., Hyman, A.A., and Rosen, M.K. (2017). Biomolecular condensates: organizers of cellular biochemistry. Nat Rev Mol Cell Biol 18, 285–298. 10.1038/nrm.2017.7.

27. Ma, H., Qi, F., Ji, L., Xie, S., Ran, J., Liu, M., Gao, J., and Zhou, J. (2022). NuMA forms condensates through phase separation to drive spindle pole assembly. J Mol Cell Biol 14. 10.1093/jmcb/mjab081.

28. Sun, M., Jia, M., Ren, H., Yang, B., Chi, W., Xin, G., Jiang, Q., and Zhang, C. (2021). NuMA regulates mitotic spindle assembly, structural dynamics and function via phase separation. Nat Commun 12, 7157. 10.1038/s41467-021-27528-6.

29. Finegan, T.M., Deem, K., Tan, R., Lowe, N., Weeks, N., Linhoff, M.W., Wang, X., López-Molini, I., Benraiss, A., and Bergstralh, D.T. (2025). Alternative splicing repurposes the Drosophila mitotic regulator Mud for meiotic functions. Genetics 231. 10.1093/genetics/iyaf145.

30. Zhu, J., Wen, W., Zheng, Z., Shang, Y., Wei, Z., Xiao, Z., Pan, Z., Du, Q., Wang, W., and Zhang, M. (2011). LGN/mInsc and LGN/NuMA complex structures suggest distinct functions in asymmetric cell division for the Par3/mInsc/LGN and Gαi/LGN/NuMA pathways. Mol Cell 43, 418–431. 10.1016/j.molcel.2011.07.011.

31. Wake, N., Weng, S.L., Zheng, T., Wang, S.H., Kirilenko, V., Mittal, J., and Fawzi, N.L. (2025). Expanding the molecular grammar of polar residues and arginine in FUS phase separation. Nat Chem Biol 21, 1076–1088. 10.1038/s41589-024-01828-6.

32. Wang, J., Choi, J.M., Holehouse, A.S., Lee, H.O., Zhang, X., Jahnel, M., Maharana, S., Lemaitre, R., Pozniakovsky, A., Drechsel, D., et al. (2018). A Molecular Grammar Governing the Driving Forces for Phase Separation of Prion-like RNA Binding Proteins. Cell 174, 688–699.e616. 10.1016/j.cell.2018.06.006.

33. Case, L.B., Zhang, X., Ditlev, J.A., and Rosen, M.K. (2019). Stoichiometry controls activity of phase-separated clusters of actin signaling proteins. Science 363, 1093–1097. 10.1126/science.aau6313.

34. So, C., Seres, K.B., Steyer, A.M., Mönnich, E., Clift, D., Pejkovska, A., Möbius, W., and Schuh, M. (2019). A liquid-like spindle domain promotes acentrosomal spindle assembly in mammalian oocytes. Science 364. 10.1126/science.aat9557.

35. Anzai, R., Mabuchi, A., and Hata, S. (2026). Coiled-coils as emerging drivers of liquid-liquid phase separation. J Biochem 179, 87–97. 10.1093/jb/mvaf065.

36. Frey, S., Richter, R.P., and Görlich, D. (2006). FG-rich repeats of nuclear pore proteins form a three-dimensional meshwork with hydrogel-like properties. Science 314, 815–817. 10.1126/science.1132516.

37. Gao, Y., Skowyra, M.L., Feng, P., and Rapoport, T.A. (2022). Protein import into peroxisomes occurs through a nuclear pore-like phase. Science 378, eadf3971. 10.1126/science.adf3971.

38. Mout, R., Bretherton, R.C., Decarreau, J., Lee, S., Gregorio, N., Edman, N.I., Ahlrichs, M., Hsia, Y., Sahtoe, D.D., Ueda, G., et al. (2024). De novo design of modular protein hydrogels with programmable intra- and extracellular viscoelasticity. Proc Natl Acad Sci U S A 121, e2309457121. 10.1073/pnas.2309457121.

39. Harmon, T.S., Holehouse, A.S., Rosen, M.K., and Pappu, R.V. (2017). Intrinsically disordered linkers determine the interplay between phase separation and gelation in multivalent proteins. Elife 6. 10.7554/eLife.30294.

40. Li, P., Banjade, S., Cheng, H.C., Kim, S., Chen, B., Guo, L., Llaguno, M., Hollingsworth, J.V., King, D.S., Banani, S.F., et al. (2012). Phase transitions in the assembly of multivalent signalling proteins. Nature 483, 336–340. 10.1038/nature10879.

41. Mateju, D., Franzmann, T.M., Patel, A., Kopach, A., Boczek, E.E., Maharana, S., Lee, H.O., Carra, S., Hyman, A.A., and Alberti, S. (2017). An aberrant phase transition of stress granules triggered by misfolded protein and prevented by chaperone function. Embo j 36, 1669–1687. 10.15252/embj.201695957.

42. Patel, A., Lee, H.O., Jawerth, L., Maharana, S., Jahnel, M., Hein, M.Y., Stoynov, S., Mahamid, J., Saha, S., Franzmann, T.M., et al. (2015). A Liquid-to-Solid Phase Transition of the ALS Protein FUS Accelerated by Disease Mutation. Cell 162, 1066–1077. 10.1016/j.cell.2015.07.047.

43. Eot-Houllier, G., Venoux, M., Vidal-Eychenié, S., Hoang, M.T., Giorgi, D., and Rouquier, S. (2010). Plk1 regulates both ASAP localization and its role in spindle pole integrity. J Biol Chem 285, 29556–29568. 10.1074/jbc.M110.144220.

44. Keder, A., Rives-Quinto, N., Aerne, B.L., Franco, M., Tapon, N., and Carmena, A. (2015). The hippo pathway core cassette regulates asymmetric cell division. Curr Biol 25, 2739–2750. 10.1016/j.cub.2015.08.064.

45. Sana, S., Keshri, R., Rajeevan, A., Kapoor, S., and Kotak, S. (2018). Plk1 regulates spindle orientation by phosphorylating NuMA in human cells. Life Sci Alliance 1, e201800223. 10.26508/lsa.201800223.

46. Sumara, I., Giménez-Abián, J.F., Gerlich, D., Hirota, T., Kraft, C., de la Torre, C., Ellenberg, J., and Peters, J.M. (2004). Roles of polo-like kinase 1 in the assembly of functional mitotic spindles. Curr Biol 14, 1712–1722. 10.1016/j.cub.2004.09.049.

47. Rajeevan, A., Olakkal, V., Balakrishnan, M., Chakrabarty, D., Charon, F., Noordermeer, D., and Kotak, S. (2025). Aurora A regulates the material property of spindle poles to orchestrate nuclear organization at mitotic exit. Embo j 44, 6797–6831. 10.1038/s44318-025-00564-4.

48. Ali, A., and Stukenberg, P.T. (2023). Aurora kinases: Generators of spatial control during mitosis. Front Cell Dev Biol 11, 1139367. 10.3389/fcell.2023.1139367.

49. Lee, C.Y., Andersen, R.O., Cabernard, C., Manning, L., Tran, K.D., Lanskey, M.J., Bashirullah, A., and Doe, C.Q. (2006). Drosophila Aurora-A kinase inhibits neuroblast self-renewal by regulating aPKC/Numb cortical polarity and spindle orientation. Genes Dev 20, 3464–3474. 10.1101/gad.1489406.

50. Wang, H., Somers, G.W., Bashirullah, A., Heberlein, U., Yu, F., and Chia, W. (2006). Aurora-A acts as a tumor suppressor and regulates self-renewal of Drosophila neuroblasts. Genes Dev 20, 3453–3463. 10.1101/gad.1487506.

51. Cho, N.H., Aslan, M., Taheri, A., Yildiz, A., and Dumont, S. (2025). NuMA mechanically reinforces the spindle independently of its partner dynein. Curr Biol 35, 4084–4095.e4085. 10.1016/j.cub.2025.07.028.

52. Colombo, S., Michel, C., Speroni, S., Ruhnow, F., Gili, M., Brito, C., and Surrey, T. (2025). NuMA is a mitotic adaptor protein that activates dynein and connects it to microtubule minus ends. J Cell Biol 224. 10.1083/jcb.202408118.

53. Okumura, M., Natsume, T., Kanemaki, M.T., and Kiyomitsu, T. (2018). Dynein-Dynactin-NuMA clusters generate cortical spindle-pulling forces as a multi-arm ensemble. Elife 7. 10.7554/eLife.36559.

54. Rao, L., Liu, X., Arnold, M., Okada, K., McKenney, R.J., Stengel, K., Sidoli, S., Berger, F., and Gennerich, A. (2026). Adaptor-mediated recruitment of three dyneins to dynactin enhances force generation. Nat Cell Biol. 10.1038/s41556-026-01877-0.

55. Volkov, V.A., and Akhmanova, A. (2024). Phase separation on microtubules: from droplet formation to cellular function? Trends Cell Biol 34, 18–30. 10.1016/j.tcb.2023.06.004.

56. Lu, M.S., and Prehoda, K.E. (2013). A NudE/14-3-3 pathway coordinates dynein and the kinesin Khc73 to position the mitotic spindle. Dev Cell 26, 369–380. 10.1016/j.devcel.2013.07.021.

57. Abramson, J., Adler, J., Dunger, J., Evans, R., Green, T., Pritzel, A., Ronneberger, O., Willmore, L., Ballard, A.J., Bambrick, J., et al. (2024). Accurate structure prediction of biomolecular interactions with AlphaFold 3. Nature 630, 493–500. 10.1038/s41586-024-07487-w.

58. Xu, J., and Zhang, Y. (2010). How significant is a protein structure similarity with TM-score = 0.5? Bioinformatics 26, 889–895. 10.1093/bioinformatics/btq066.

59. Zhang, Y., and Skolnick, J. (2004). Scoring function for automated assessment of protein structure template quality. Proteins 57, 702–710. 10.1002/prot.20264.

60. Madaj, R., Martinez-Goikoetxea, M., Kaminski, K., Ludwiczak, J., and Dunin-Horkawicz, S. (2025). Applicability of AlphaFold2 in the modeling of dimeric, trimeric, and tetrameric coiled-coil domains. Protein Sci 34, e5244. 10.1002/pro.5244.

