## Supplementary figures and images for "Liquid-solid phase transitions in the biological condensates of a conserved mitotic spindle regulator"

### Figure S1

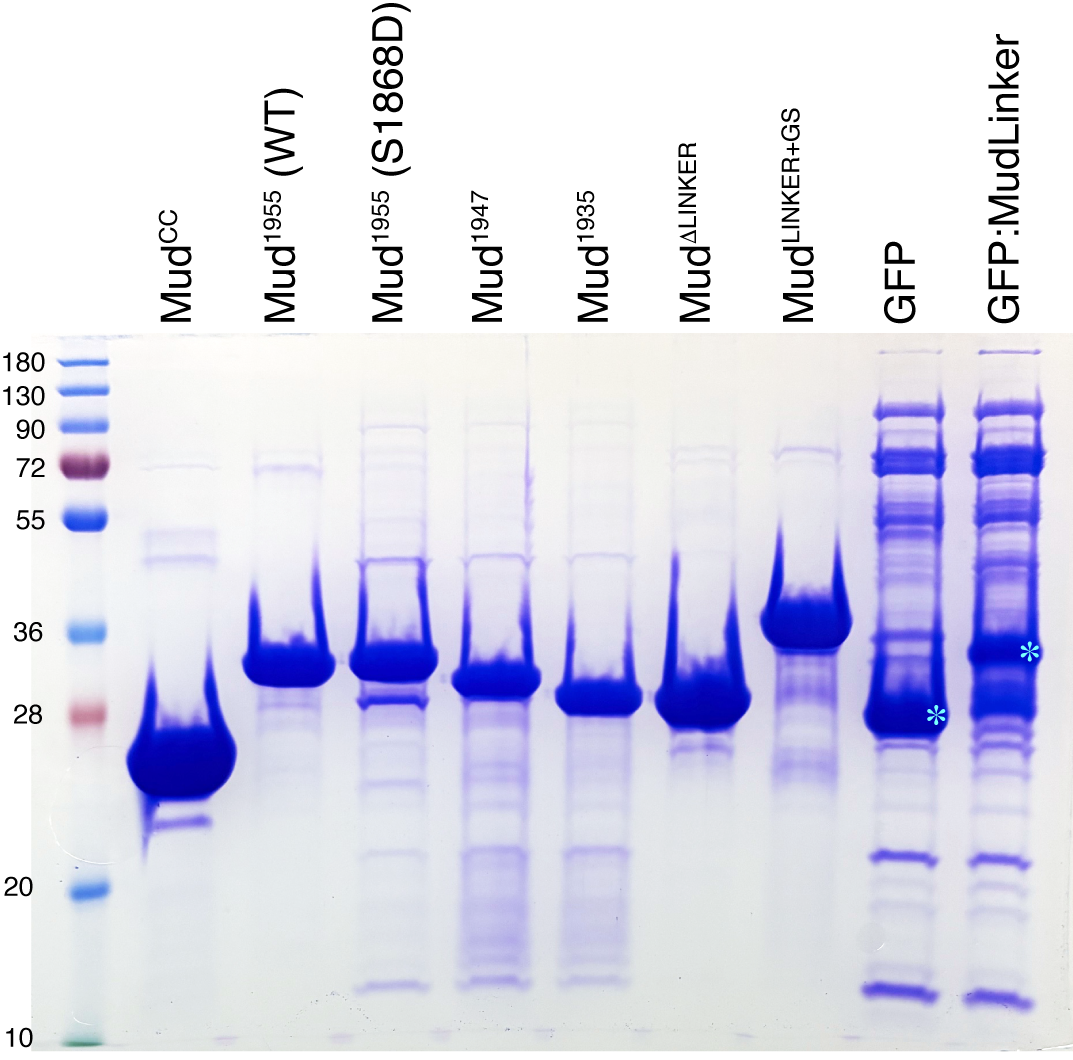

### Figure S2

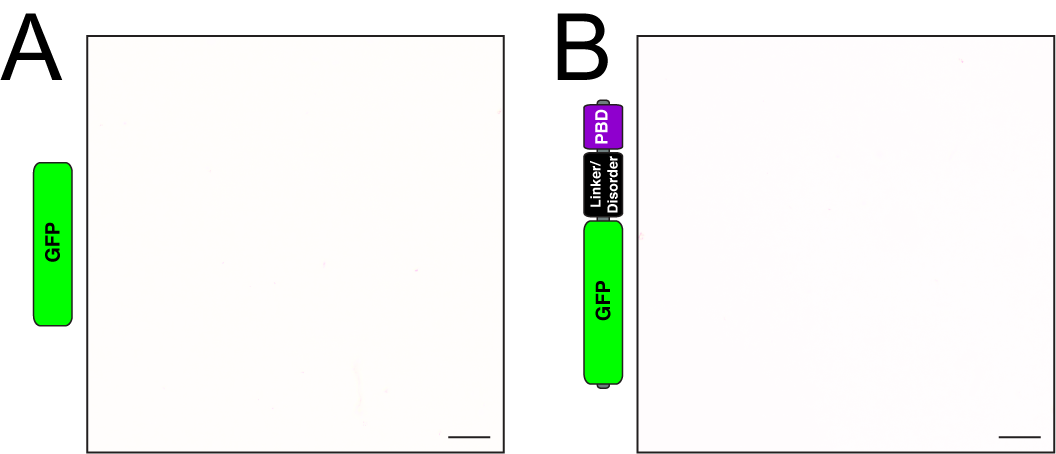

### Figure S3

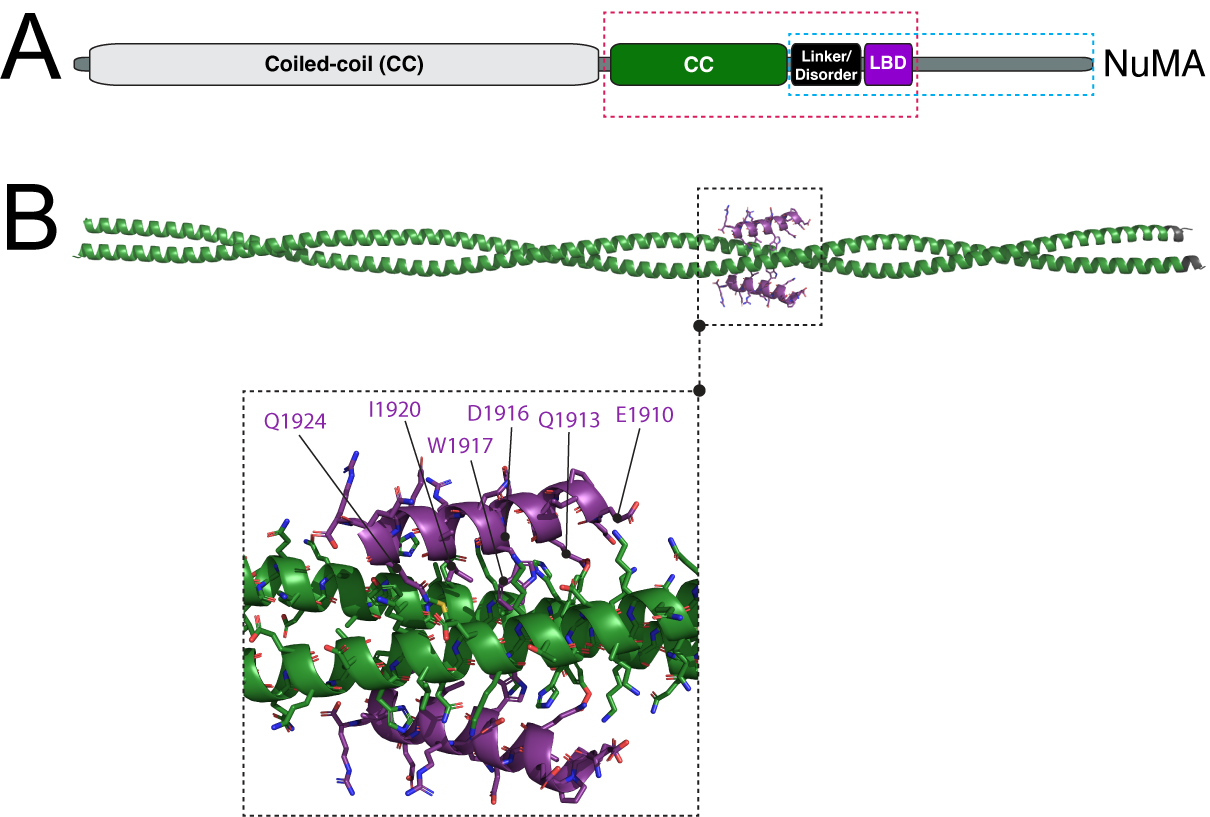

### Figure S4

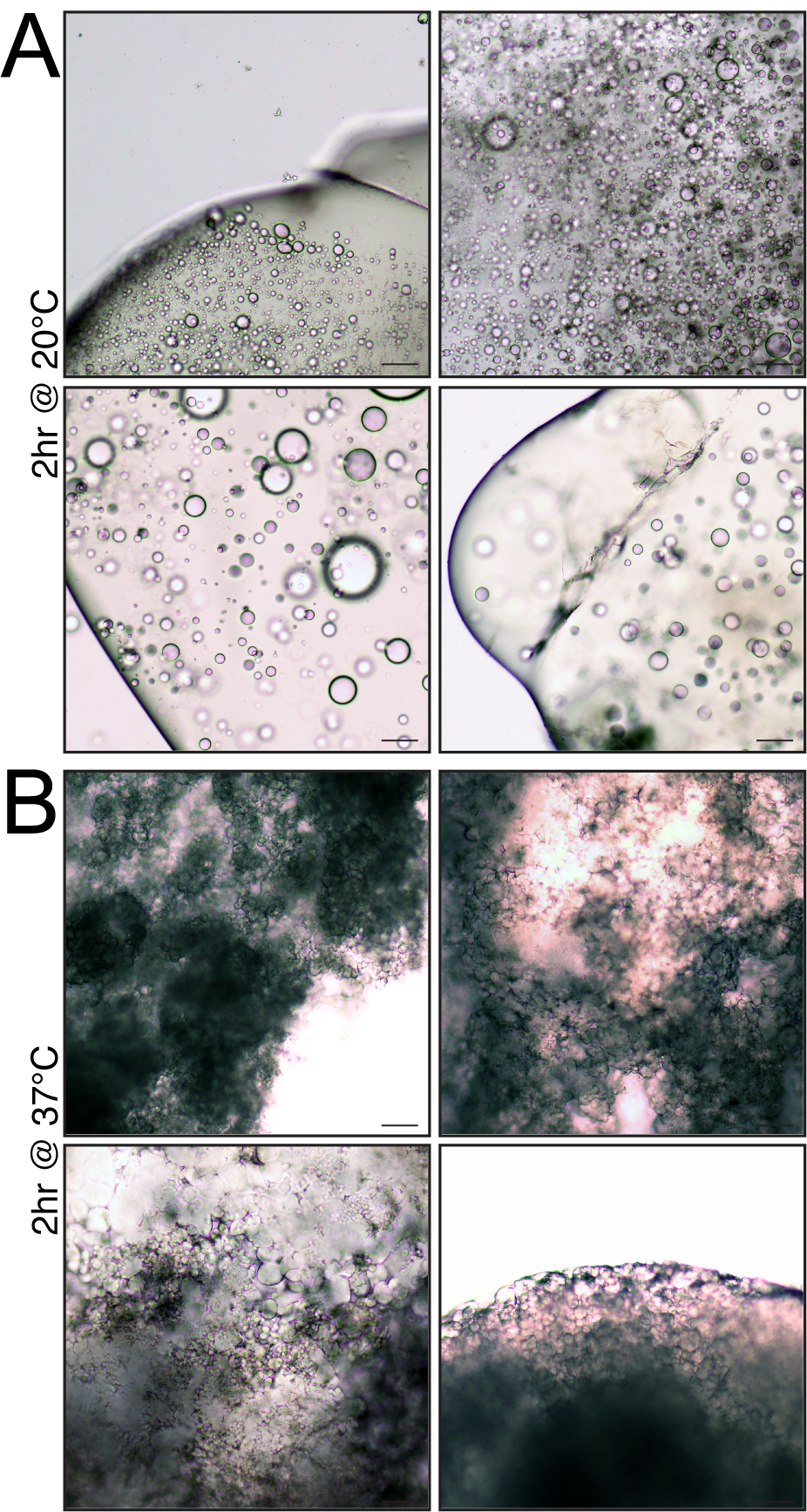
